# Glucose Challenge Uncovers Temporal Fungibility of Metabolic Homeostasis over a day:night cycle

**DOI:** 10.1101/2023.10.30.564837

**Authors:** Dania M. Malik, Seth D. Rhoades, Shirley L. Zhang, Arjun Sengupta, Annika Barber, Paula Haynes, Erna Sif Arnadottir, Allan Pack, Richard G. Kibbey, Pinky Kain, Amita Sehgal, Aalim M. Weljie

## Abstract

Rhythmicity is a cornerstone of behavioral and biological processes, especially metabolism, yet the mechanisms behind metabolite cycling remain elusive. This study uncovers a robust oscillation in key metabolite pathways downstream of glucose in humans. A purpose-built ^13^C_6_-glucose isotope tracing platform was used to sample *Drosophila* every 4h and probe these pathways, revealing a striking peak in biosynthesis shortly after lights-on in wild-type flies. A hyperactive mutant (*fumin*) demonstrates increased Krebs cycle labelling and dawn-specific glycolysis labelling. Surprisingly, neither underlying feeding rhythms nor the presence of food availability explain the rhythmicity of glucose processing across genotypes, suggesting a robust internal mechanism for metabolic control of glucose processing. These results align with clinical data highlighting detrimental effects of mistimed energy intake. Our approach offers a unique insight into the dynamic range of daily metabolic processing and provides a mechanistic foundation for exploring circadian metabolic homeostasis in disease contexts.

## Introduction

Rhythmicity is an ever-present feature of physiology driven by zeitgebers (environmental cues), including light and temperature, that entrain endogenous circadian clocks. Molecular circadian clocks coordinate biological rhythmicity within each tissue, while modern pressures such as mistimed light exposure, shift work, and social jetlag disrupt the evolved circadian system^1–3^. Mistimed clocks play a role for increasing risk of diseases including obesity, cardiovascular disease, and cancer^4–8^. Recent molecular studies of human biofluids have shown strong rhythmicity of transcripts and metabolite levels under strict laboratory-controlled conditions^9,10^. In reality, humans vary in chronotype and display inter-individual variation in the number of rhythmic compounds and related measures (phases and amplitudes)^11–14^. In spite of this variation, certain metabolic phenomena, such as the diurnal response to nutrient intake, are relatively robust and reproducible. Glucose-related measurements such as glucose tolerance and post-feeding have shown time variation with higher sugar levels reported in the afternoon versus morning measurements^15–17^ (so-called ‘afternoon diabetes’). Moreover, afternoon measurements have been noted to allow or exclude diagnoses of diabetes where morning measurements are borderline^15^. In spite of this clinical significance, the mechanistic pathways downstream of glucose metabolism which underlie these temporal differences in glucose processing remain largely unknown.

At the molecular level, model organisms play an important role in the study of metabolic rhythmicity^18–21^. They provide the advantage of providing isogenic genetic backgrounds for study and a number of studies probing clock protein function^22^ and zeitgebers such as light and feeding ^23–25^ have provided valuable insight into steady state metabolic pool changes at specific points in time. Our recent study^26^ using an ion-switching LC-MS.MS method has shown altered metabolism during the dark period in *Drosophila* short sleep mutants. However, since metabolism occurs in a network of interacting pathways, changes in the level of a given metabolite can arise from multiple possibilities and mechanistic interpretation is, by necessity, only inferred in studies where only metabolite pools are measured. Metabolic tracing provides a direct avenue to address the mechanism by which altered metabolite levels arise^27^. To date, information on diurnal rhythmic metabolic function via tracing is remarkedly sparse. Isotope tracer studies are used for clinical studies of glucose metabolism, but the fate of metabolites downstream is not known. We have recently demonstrated rhythmicity of pool sizes for glucose and related metabolites (glucose-6-phosphate and ribose-5-phosphate) in human red blood cells^28^. The further use of labelled glucose allowed for insight into rhythmicity driven specifically by glycolytic pathways and the pentose phosphate pathway noting opposite phases^28^. Other studies have also used stable and/or radioactive isotope tracers in conjunction with targeted approaches in circadian contexts; however, limited number of time points have been measured^29–32^ so while important insight into some time-dependent differences can be ascertained, higher time resolution data are needed for reliable detection of rhythmicity^18,33^.

In this study, we first examined the time-of-day variability of metabolite pools in healthy humans to characterize the rhythmic activity of pathways downstream of glucose metabolism. Based on this set of pathways, which are largely conserved across species, we developed a customized analytical platform to identify physiological metabolite homeostatic rhythms in *Drosophila* via isotope tracing of glucose. The use of model organisms with isotope labelled metabolites not only allows for greater flexibility in experimental designs, but also allows us to establish a guide for eventual clinical studies. Toward this goal, we probed the circadian processing of a glucose bolus in *Drosophila* as its core clock and many of the glucose processing and insulin signaling pathways are conserved in mammals^34–36^. A number of rhythmic downstream glucose products were observed in both WT Drosophila as well as a hyperactive mutant, *fumin* (*fmn*)^37^, with peaks in biosynthesis from glucose during the light phase which provides direct evidence for glucose processing limitation later in the day. More generally, the platform allows for metabolic tracing through the introduction of a small volume of an isotope tracer and can be adapted for use in other model organisms or humans to study isotope fate rhythmicity with high time resolution.

## Results

### Total metabolite pools reveal time of day variation in glucose-related metabolites in humans

Glucose metabolism is tightly regulated by circadian processes and disruption results in various disorders, thus we first looked to establish the extent and robustness of time-of-day variability in a network of glucose-related metabolites using steady state measurements in humans. A curated set of 93 metabolites were measured in plasma samples collected every 4 hours over 24 hours (N = 14 subjects; Extended Data Fig. 1a) using a targeted ion-switching LCMS method^38^. Multivariate analysis of the resulting data exhibits remarkably robust time-of-day variability (Fig. 1a). Amino acids and related metabolites such as alanine, ornithine, cystathionine, and asparagine were noted to be elevated around ZT 8 and ZT 12 while elevated levels of TCA cycle metabolites such as citrate/isocitrate and malate were associated with ZT 0 and 4 (Fig. 1b). This result implies that the total pools of metabolites are dramatically altered throughout the day, hinting at differential pathways being available for nutrient processing. Pairwise multivariate models distinguished each single time point from a combination of the others (‘one vs all’) with the exception of ZT 4. (Fig. 1c and Extended Data Fig. 1b to f). Pathway analysis of each time-point model using significant metabolites demonstrates a unique set of pathways enriched at each specific time of day. For example: alanine, aspartate and glutamate metabolism and cysteine and methionine metabolism were significantly different during the dark period; TCA cycle and valine, leucine, and isoleucine biosynthesis were significantly different throughout the day; phenylalanine, tyrosine, and tryptophan biosynthesis were significant across all of the time points, but not at a specific one (Fig. 1d). Taken together the multivariate approach indicated overall differences in metabolite levels and activity of pathways at different time of the day.

**Fig. 1:**
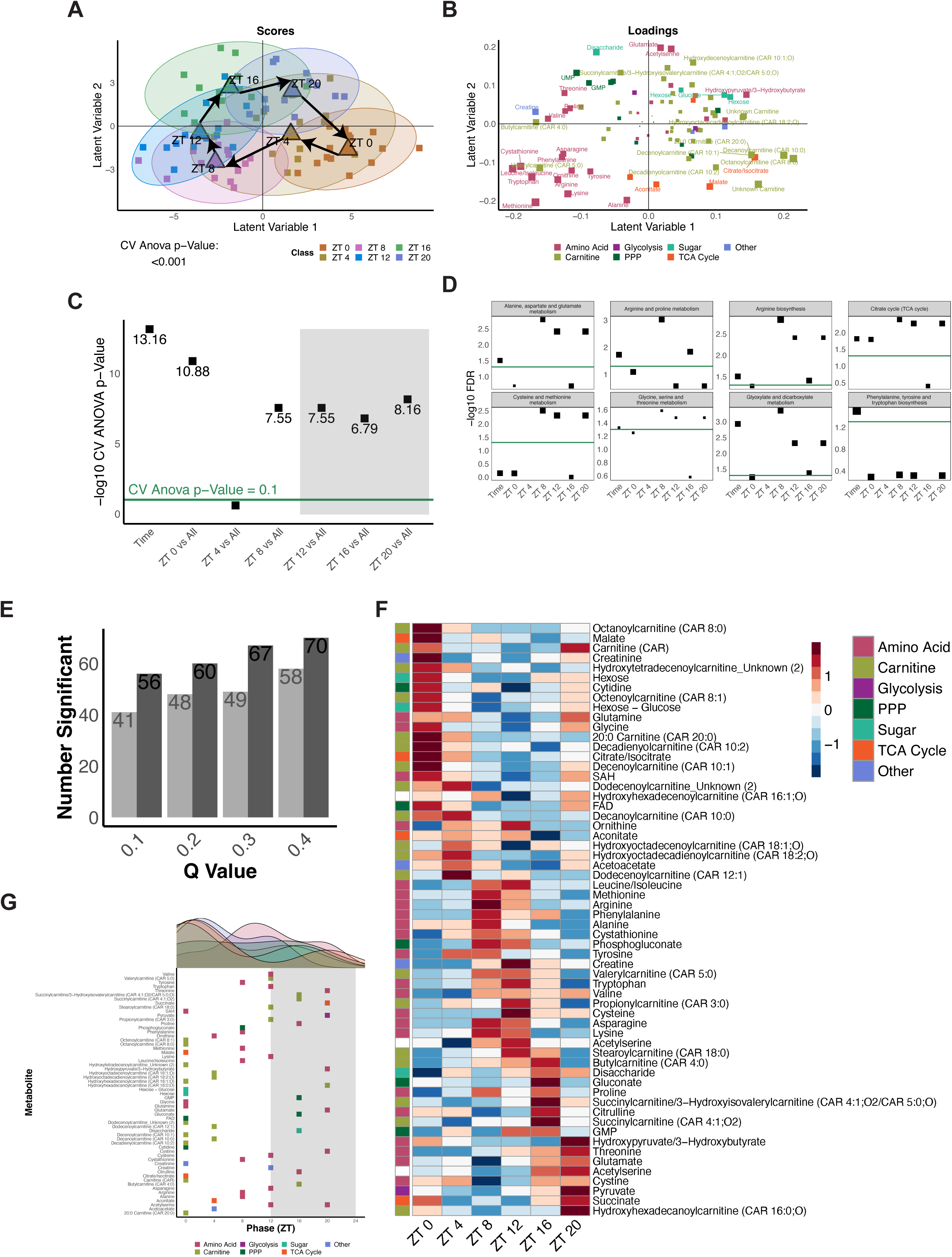
Steady state metabolite levels in human plasma samples reveal time of day variation in glucose-related metabolites. **a,** Scores plot for the significant Time OPLS-DA Model. Classes were defined as time points (ZT 0, ZT 4, ZT 8, ZT 12, ZT 16, and ZT 20). Colors represent classes (ZT times), squares represent individual samples and triangles represent the group average. **b,** Corresponding loadings plot for the Time OPLS-DA Model. Colors represent classes of metabolites and size represents significance as defined through VIP values. **c,** Negative log CV-ANOVA p-values for overall time and each time point tested in pairwise OPLS-DA models. Significance was defined as a CV-ANOVA p-value less than 0.1 (shown as points above the green line). **d,** Negative log FDR values for pathways significant in at least one significant OPLS-DA model (defined in c). Size represents impact and significance was defined as a FDR value less than 0.05. **e,** Overview of the number of significant 24-hr rhythmic metabolites observed at RAIN (dark) or JTK (light) q-value cut-offs of 0.1, 0.2, 0.3, and 0.4. **f,** Phase ordered heatmap of significantly cycling metabolites with 24-hr periods as tested by RAIN with a q-value less than 0.2. **g,** Distribution of RAIN phases for significant 24-hr metabolites. Colors represent classes of metabolites.

Formal rhythmicity testing was then conducted and 24-hr and 20-28-hr rhythms were observed across different q-value cut-offs (q<0.2, 03, or 0.4, Fig. 1e and Extended Data Fig. 1g). Metabolites with a RAIN q-value less than 0.2 were selected for further analysis (Fig. 1f and Extended Data Fig. 1h). Phases of cycling metabolites were found to be distributed across the day with a peak observed during the early light period (ZT 0 to 4) (Fig. 1g and Extended Data Fig. 1i). Taken together, these results imply that glucose tolerance is influenced by differential downstream pathways regulated in a time of day dependent manner. Notably the robustness of the response and similarity across the 14 subjects was remarkable and not obvious from individual metabolites levels. This suggests that the concerted network of glucose metabolism provides constraints under which observations of glucose metabolism should be contextualized.

### Overview and validation of isotope tracing platform

The pathways measured related to glucose metabolism are largely conserved across many species, and thus we developed a mass spectrometry-based platform to trace the fate of glucose metabolites through a broad network of downstream metabolism as observed in the human study above for testing in *Drosophila*. The use of stable isotope tracers such as glucose enable the metabolites of interest to be monitored at the atomic level, thus allowing further insights into how glycemic metabolism is altered as a function of time. Uniformly labelled ^13^C_6_-glucose was chosen as the stable isotope tracer as it allows carbon flow to be monitored into downstream metabolites spanning the TCA cycle, pentose phosphate pathway (PPP), glycolysis, and amino acid biosynthesis. A range of glucose concentrations were tested in *Drosophila* to identify an optimal concentration at which *Drosophila* remained viable. Flies were sampled one hour after introducing the tracer to allow glucose to be metabolized into downstream pathways and to maintain sufficient time resolution to monitor diurnal shifts. The 1 M bolus of glucose was introduced intra-thoracically using a blue dye as a visual aid. The injections were performed on flies that were sedated with cold treatment to prevent metabolic changes caused by hypoxia due to carbon dioxide exposure (Fig. 2a and 2b). Representative images of a needle containing the injection solution highlight the small volume (∼31 nl/fly) introduced into flies (Fig. 2b). Since a non-physiological glucose concentration (∼25.6 µM/fly corresponding to ∼0.8 pmol/fly or ∼148 pg/fly) was used to allow for detection by mass spectrometry, geotaxis and locomotion assays were used to assess whether the glucose and/or blue dye were impacting physiology. No significant differences were noted between the glucose and PBS injected conditions for both climbing ability and activity (Fig. 2c and d). This indicated that the supraphysiological glucose and dye concentrations did not impair geotaxis or locomotion of WT flies and thus were chosen as a tracer to assess time resolved metabolic nutrient challenge.

**Fig. 2:**
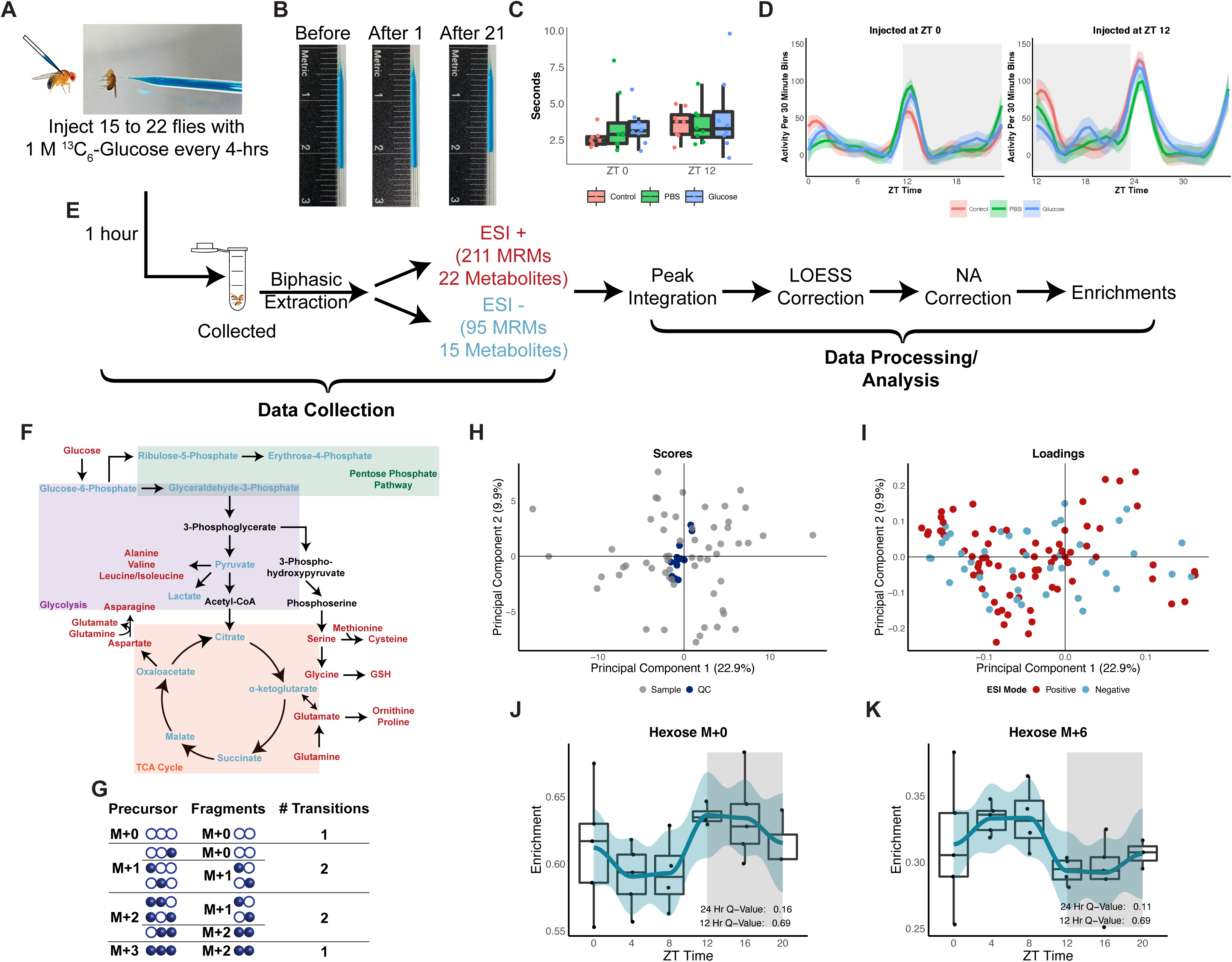
Overview and validation of isotope tracing platform and processing methods of flies injected with a labeled bolus of glucose. **a,** Image of a fly being injected with a glucose and blue dye solution. **b,** Representative image of a needle containing a 1 M Glucose and 0.504 mM FCF solution before injections (left), after injecting 1 fly (middle) and after injecting 21 flies (right). **c,** climbing ability of male flies assessed through a geotaxis assay as the time needed to climb 4 cm after no injections (control) and injections with 1 M glucose or PBS. Two-sided t-test was used with significance defined as a p-value less than 0.05. n = 5 to 8 per group and time point. **d,** Locomotion assay of male flies without injections (control) or after injections with 1 M Glucose or PBS at ZT 0 (left panel) or at ZT 12 (right panel). Activity was recorded for 24 hours after injections. LOESS trace is shown for activity counts for each 30-minute time period. Significance was tested using a two-sided t-test for each activity bin across groups with significance defined as a BH adjusted p-value (q-value) less than 0.05. n = 8 per group and time point. **e,** Simplified overview of workflow. Flies (n = 15 to 23) injected with 1 M Glucose were collected 1 hour post injection. Upper fractions (polar metabolites) collected from a biphasic extraction were analyzed in positive and negative electrospray ionization modes. Acquired data was integrated to obtain ion counts which were used for an instrumental drift correction and normalized to the amount of blue dye injected. The normalized data is natural abundance corrected and used to calculate enrichments for further analysis. **f,** Simplified view of downstream glucose metabolism with metabolites included in MS methods highlighted in red (ESI+) and blue (ESI-). Shading indicates downstream glucose pathways. **g,** Depiction of labelling patterns considered for generating MRM transitions of a 3-carbon compound, such as alanine, with a 2-carbon fragment. In generating transitions, the number of potential labelled carbons in the precursor and fragments are considered for each isotopologue. **h,** PCA scores plot of biological (light grey) and QC samples (blue) showing clustering of the QC samples. **i,** PCA loadings plot corresponding to the scores plot (h) showing the distribution of metabolite enrichments detected in positive (red) and negative (blue) modes. **j, k,** Box plot of enrichments along with a LOESS trace of hexose M+0 (**j**) and hexose M+6 (**k**) observed across a day at 4 hr intervals in wild type flies. n = 4 to 5 biological replicates of 15 male flies per time point. Rhythmicity of 24-and 12-hr periods was tested using RAIN and significance was defined as a BH adjusted q-value less than 0.2.

We challenged WT *Drosophila* with 1 M uniformly labelled ^13^C_6_-glucose at 4-hour intervals across a day to determine how glucose is processed at different times in the fly body (Fig. 2e). Each sample was analyzed using two LCMS methods covering 22 and 15 downstream glucose metabolites in electrospray ionization positive (211 transitions) and negative (95 transitions) modes, respectively (Fig. 2e and f). Metabolite transitions were defined by considering the number of carbons and possible labelling patterns of the precursor and fragment (Fig. 2g). The resulting data were corrected for instrumental drift and normalized to the blue dye ion counts to adjust for variability in injection volumes before performing a natural abundance (NA) correction. Enrichments were calculated using the NA corrected data and were used for further analyses (Fig. 2e).

To determine data quality, enrichments for all biological and quality control (QC) samples were visualized using a PCA (Fig. 2h and i). Overall analytical quality was robust as noted through clustering of the QC samples in the scores plot (Fig. 2h) and evenly distributed compounds from both ionization modes in the loadings plot (Fig. 2i). Furthermore, the platform was sensitive to detect changes in levels of detected isotopologues one hour after samples were challenged with labelled glucose at 4-hour intervals. For example, opposing changes were noted in the enrichments of two hexose isotopologues, M+0 and M+6 (Fig. 2j and k). Here, hexose M+0 (composed primarily of glucose) peaked around ZT 12 (lights off) while the M+6 (composed of all labelled glucose) form peaked around ZT 4. This points to potential changes in overall glucose usage during the day.

### Increased biosynthesis from glucose challenge is distributed across multiple pathways at ZT 4

To further understand changes in glucose utilization as a function of time in WT Drosophila, JTK and RAIN algorithms were used to test for 24-, 20-28-, and 12-hr rhythmicity. As the FDR (q-value) cut-off was increased from 0.1 to 0.4, increasing numbers of isotopologues cycled with periods of 24-, 20-28-, and 12-hr periods (Fig. 3a, Extended Data Fig. 2a and d). For further analyses, rhythmic compounds with a FDR less than 0.2 were used (Fig. 3b, Extended Data Fig. 2b and e). For all periods tested, metabolite pools (sum of all isotopologues for a given metabolite) were not significantly rhythmic. For 24-and 20-28-hr rhythmic results, labelled isotopologues peaked at ZT 4 while unlabeled (M+0) isotopologues peaked at the light: dark (LD) transition (ZT 12) (Fig. 3c and Extended Data Fig. 2c) which is to be expected based on the hexose labelling pattern. On the other hand, 12-hr rhythmic isotopologues peaked mostly around ZT 8 and ZT 20 (Extended Data Fig. 2f). Since 24-hr rhythmicity was not present in metabolite pools, this suggests labelled glucose carbons were incorporated into isotopologues such as serine M+3, proline M+2, M+3, M+4, and M+5 at ZT 4. Taken together, peaks at ZT 4 for erythrose-4-phosphate M+4 and AMP M+5 can imply use of the PPP while M+3 forms of serine and alanine may arise downstream of glycolysis, and glutamine and proline isotopologues from the TCA cycle (Fig. 3d). This points to overall increased activity of downstream glucose metabolic pathways uniquely at ZT 4, which we will refer to as a ‘rush hour’ of biosynthesis.

**Fig. 3:**
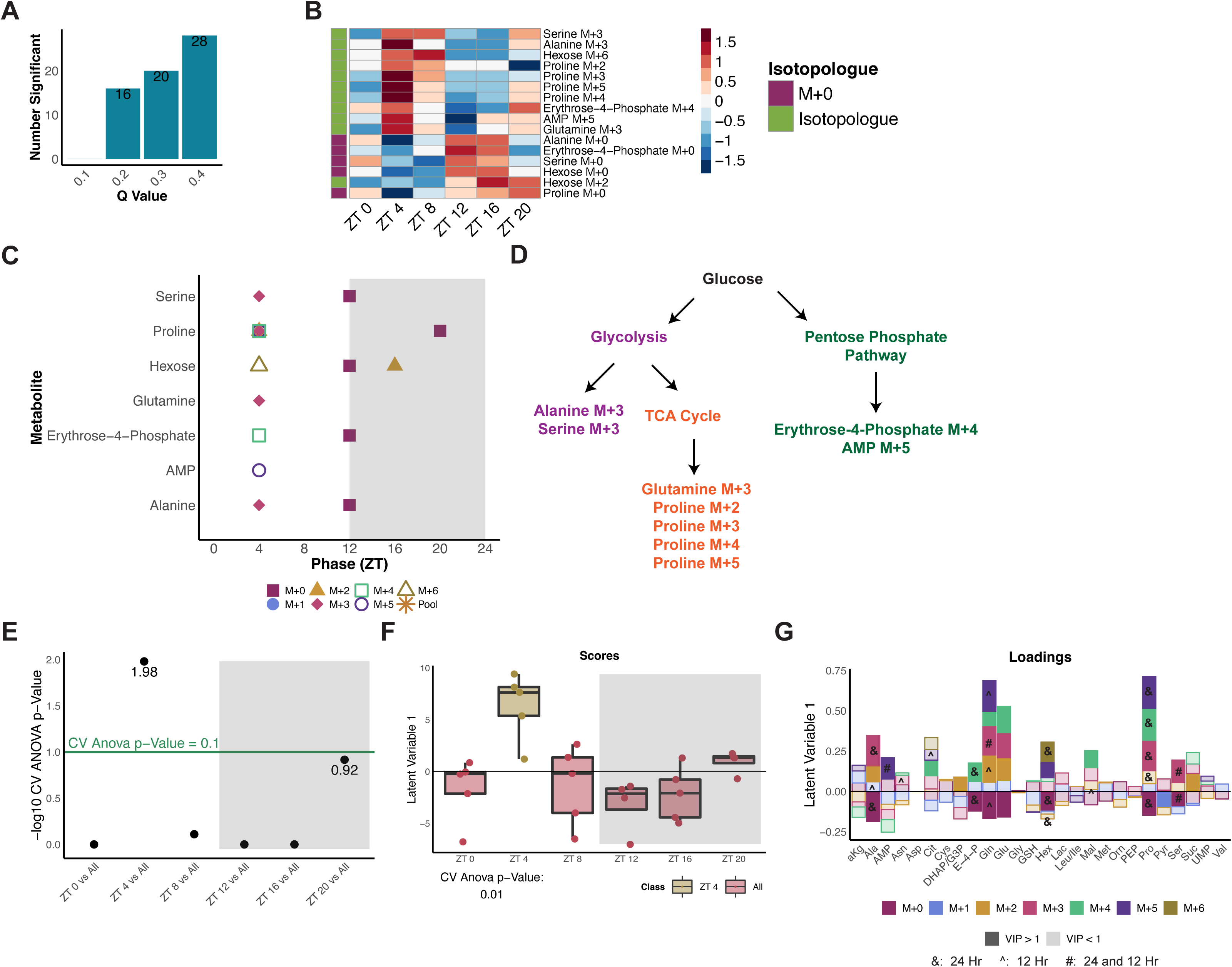
Increased biosynthesis from glucose challenge is distributed across multiple pathways at ZT 4. **a,** Overview of the number of significant 24-hr rhythmic compounds observed at RAIN q-values of 0.1, 0.2, 0.3, and 0.4. **b,** Phase ordered heatmaps of significantly cycling compounds with 24-hr periods in WT as tested by RAIN with a q-value less than 0.2. **c,** Distribution of RAIN phases for significant 24-hr compounds grouped by metabolite. Shapes and colors refer to isotopologues or pools. **d,** A simplified overview of downstream metabolism indicating potential routes for the biosynthesis of 24-hr rhythmic isotopologues from b. **e,** Negative log CV-ANOVA p-values for each time point tested in pairwise OPLS-DA models. Significance was defined as a CV-ANOVA p-value less than 0.1 (shown as points above the green line. **f,** Scores plot for the significant WT ZT 4 vs All OPLS-DA Model. Two classes were defined as ZT 4 and All (ZT 0, ZT 8, ZT 12, ZT 16 and ZT 20). **g,** Corresponding loadings plot for the WT ZT 4 vs All OPLS-DA model. Colors represent isotopologues with shading representing VIP significance (dark signifies a VIP value greater than 1 (significantly contributing to separation of ZT 4 from other time points) while lighter shades are used for a VIP less than 1). Rhythmic isotopologues from a and Extended Data Fig 2e are indicated by symbols (& refers to significant 24-hr rhythmicity only, ^ refers to significant 12-hr rhythmicity only, # refers to significant 24- and 12-hr rhythmicity). alpha-ketoglutarate (a-kG); alanine (Ala); adenine monophosphate (AMP); asparagine (Asn); aspartate (Asp); citrate (Cit); cysteine (Cys); dihydroxyacetone phosphate (DHAP); glyceraldehyde-3-phosphate (G3P); erythrose-4-phosphate (E-4-P); glutamine (Gln); glutamate (Glu); glycine (Gly); glutathione (GSH); hexose (hex); lactate (Lac); leucine (Leu); isoleucine (Ile); malate (Mal); methionine (Met); ornithine (Orn); phosphoenolpyruvate (PEP); proline (Pro); serine (Ser); succinate (Suc); uridine monophosphate (UMP); valine (Val).

To determine if time-of-day differences in glucose usage persisted at an overall metabolic profile-level, a multivariate approach using orthogonal partial least squares-discriminant analysis (OPLS-DA) models was employed to determine whether any time point was unique from the rest. From all time points tested, ZT 4 emerged as a distinct time point compared to all remaining time points (Fig. 3e and f). Higher enrichments of isotopologues such as alanine M+2, glutamine M+2 to M+5, glutamate M+2 to M+4, and proline M+3 to M+5 and lower enrichments of M+0 forms (alanine, glutamine, glutamate, and proline) were associated with ZT 4 (Fig. 3g). This further supported the rhythmicity analysis in highlighting increased incorporation of labelled glucose into downstream products at ZT 4 suggesting increased biosynthesis.

### The TCA Cycle is preferentially used in a hyperactive mutant in response to a glucose challenge

As activity levels can act as a weak zeitgeber to influence the clock^39^, we next sought to determine how glucose is used differently in a hyperactive short-sleeping mutant deficient in dopamine reuptake, *fumin* (*fmn*)^37^. Overall, *fmn* displays elevated activity profiles as compared to WT with higher levels observed during the dark period (Fig. 4a). We challenged the mutant with glucose in a similar manner to WT. The climbing ability and activity of *fmn* were also not impaired in flies injected with glucose vs PBS at both ZT 0 and ZT 12 (Fig. 4b and c). Therefore, the injection of supraphysiological concentrations of glucose did not dramatically alter the physiology of the flies.

**Fig. 4:**
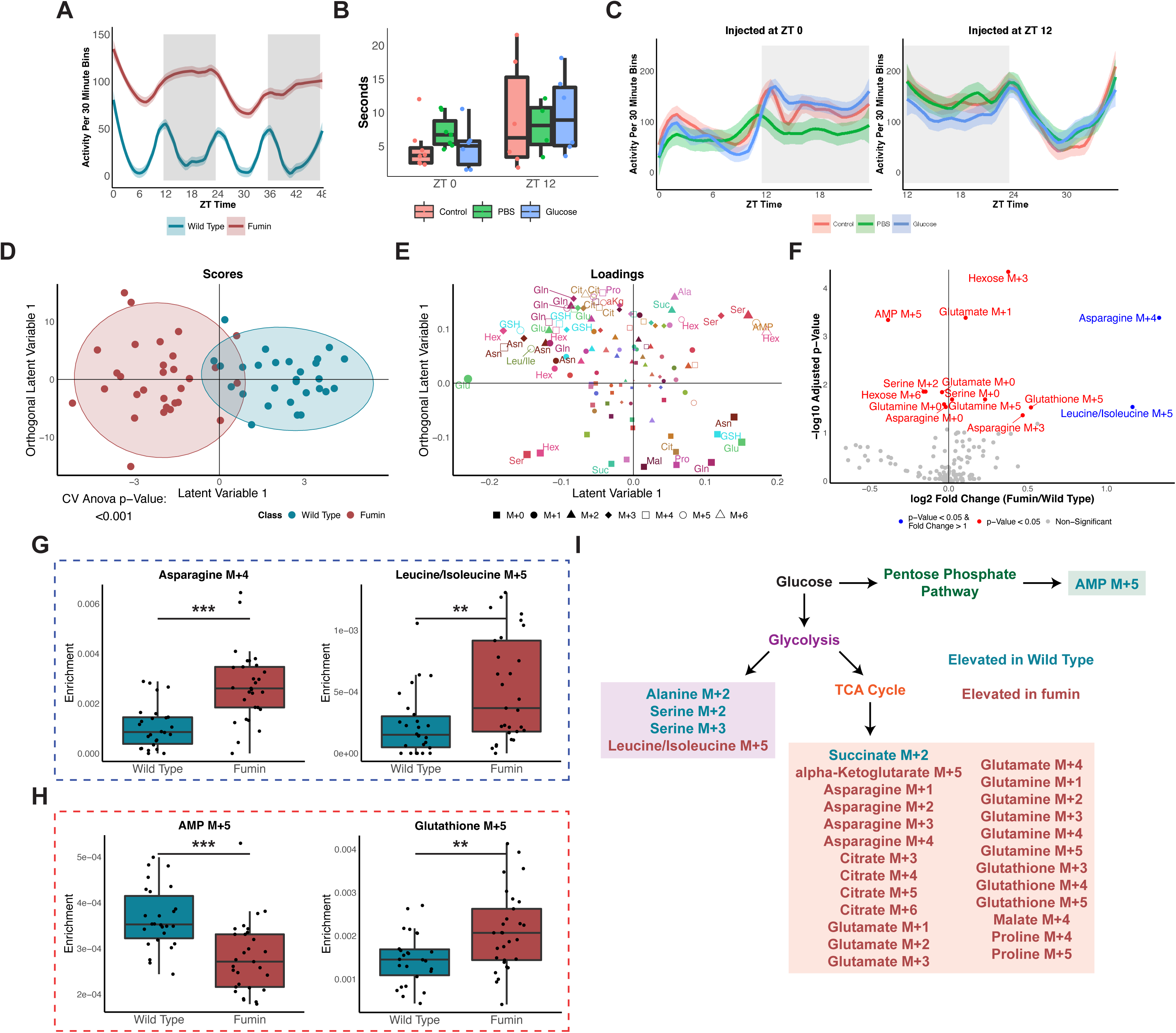
The TCA Cycle is preferentially used in a hyperactive mutant in response to a glucose challenge. **a,** LOESS trace of activity counts per 30 min bins for WT and *fmn* over 48 hr under 12-hour LD conditions. **b,** Climbing ability of male *fmn* flies assessed through a geotaxis assay as described in Fig. 2c. **c,** Locomotion assay of male *fmn* flies assessed through a locomotion assay as described in Fig. 2d. **d,** A two-component OPLS-DA scores plot of WT vs *fmn*. All time points collected (ZT 0, 4, 8, 12, 16 and 20) are included for both classes defined as genotypes. n = 4 to 5 biological replicates with 15 male flies per time point and genotype. **e,** Corresponding loadings plot for the WT vs *fmn* OPLS-DA model (d). Isotopologues are represented by shapes, metabolites by color and size of the shapes represents significance as defined through VIP values. Abbreviations as defined in Fig. 3g. **f,** Volcano plot of enrichments analyzed through univariate analyses. log2 of the fold change between *fmn* and WT is shown on the x-axis while the negative log10 BH adjusted genotype p-value from a two-way ANOVA is shown on the y-axis. A fold change cutoff of greater than 1 was used and a p-value less than 0.05 was defined as significant. **g,h,** Representative boxplots of enrichments by genotype for significant isotopologues by p-value and fold change (g) or by p-value only (h). **i,** A simplified overview of downstream glucose metabolism indicating potential routes for the biosynthesis of VIP metabolites from e. Isotopologue text color refers to the genotype with higher loadings.

Overall, wild type and *fumin* process glucose distinctly as indicated by a significant two-component discriminant (OPLS-DA) model (Fig. 4d). Hexose M+6 enrichments were relatively higher in WT while M+0 enrichments were relatively higher in *fmn* suggesting that labelled glucose was more readily used in *fmn*. Overall, enrichments of primarily labelled isotopologues (i.e., glutamate M+1, glutathione M+5, and asparagine M+2 and M+4) contributed to the separation of *fmn* from WT while enrichments of several unlabeled isotopologues such as asparagine, glutamate, and glutathione were more associated with WT samples (Fig. 4e). Univariate analyses of genotype differences also showed similar changes (Fig. 4f). For instance, enrichments of asparagine M+4 and leucine/isoleucine M+5 were significantly higher (at least 2x) in *fmn* as compared to WT (Fig. 4g). Since leucine and isoleucine are essential amino acids, the synthesis of these may be a byproduct of the fly microbiome. Additional significant differences such as higher AMP M+5 levels in WT and elevated glutathione M+5 levels in *fmn* were noted through the adjusted genotype p-value (Fig. 4h). As glutathione M+5 requires either glutamine M+5, or glutamine M+3 and glycine M+2 as precursors, we also looked at the correlation of these isotopologues. Glutathione M+5 enrichments were found to be significantly correlated with glutamine M+3 (p<0.05) and weakly correlated with glutamine M+5 (p<0.1) enrichments in *fmn* (Extended Data Fig. 3a). From both the univariate and multivariate approaches, relatively higher enrichments of AMP M+5, a metabolite related to the PPP, and metabolites associated with glycolysis (alanine M+2, serine M+2 and M+3) were observed in WT while metabolites involved in or downstream of the TCA cycle (alpha-ketoglutarate M+5, asparagine, glutamate, and glutamine isotopologues) had relatively higher enrichments in *fmn* (Fig. 4i). Overall, both multivariate and univariate approaches support increased TCA cycle labelling in *fmn* while glycolysis and PPP labelling is associated with WT.

### fumin has two biosynthetic ‘rush hours’ at dawn and dusk

In addition to overall genotype-level differences, we sought to determine how the two genotypes differed in glucose utilization through time. Through pairwise OPLS-DA models of the genotypes at each time point, a significant model was observed at ZT 4 (Extended Data Fig. 3b and c). Three types of changes were noted in labelled isotopologues: 1. Higher relative enrichments in WT (i.e., alanine M+1 to M+3 and proline M+3 to M+5); 2. Higher relative enrichments in *fmn* (i.e., alpha-ketoglutarate M+ 2 to M+4, asparagine M+1 to M+4, and glutathione M+1, M+2 and M+5); and 3. Higher enrichments spread across the two genotypes (i.e., higher AMP M+5 and lactate M+3 in WT vs higher AMP M+1 and lactate M+2 in *fmn*) (Extended Data Fig. 3d). This suggests differences in downstream glucose products and/or differences in glucose utilization pathways in WT when compared to *fmn*. Overall, both multivariate and univariate approaches supported the idea that glucose is differentially used between the two genotypes at ZT4 with increased TCA cycle labelling observed in *fumin*.

Next, to determine whether glucose is processed differently by *fumin* based on time-of-day, pairwise discriminant models were generated to test whether a time point(s) was distinct from all other time points. Both ZT 0 and ZT 12 emerged as being significantly distinct from other time points through one-and two-component models respectively (Fig. 5a, b, and d). Increased enrichments of labelled isotopologues were noted at both time points, ZT 0 and ZT 12, pointing to increased incorporation of glucose carbons into the isotopologues (Fig. 5c and e). Furthermore, the light time points (ZT 4 and 8) were significantly different from the dark ones (ZT 16 and 20) in an OPLS-DA model (Extended Data Fig. 4a). The light time points were associated with relatively increased enrichments of labelled isotopologues (Extended Data Fig. 4b).

**Fig. 5:**
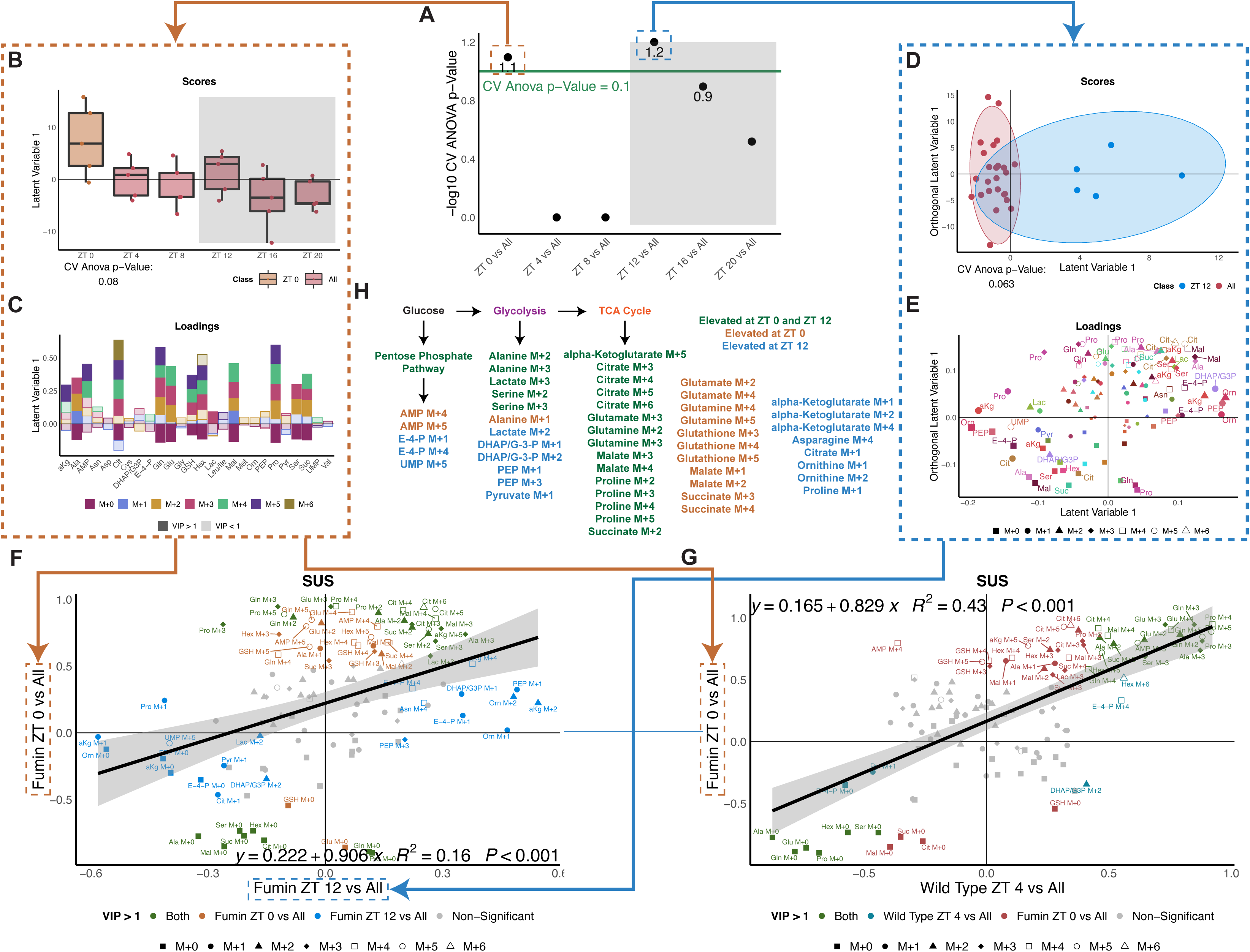
Excess glucose processing is unique at dawn and dusk compared to other times of day in *fumin*. **a,** Negative log CV-ANOVA p-values for each time point tested in pairwise OPLS-DA models for *fmn*. Significance was defined as a CV-ANOVA p-value less than 0.1 (Shown as points above the green line). **b,** Scores plot for significant *fmn* ZT 0 vs All OPLS-DA model. Two classes were defined as ZT 0 and All (ZT 4, ZT 8, ZT 12, ZT 16, ZT 20). **c,** Corresponding loadings plot for the *fmn* ZT 0 vs All OPLS-DA model. Colors represent isotopologues with shading representing VIP significance (dark signifies a VIP value greater than 1 while lighter shades are used for a VIP less than 1). Abbreviations are defined in Fig. 3g. **d,** A two-component scores plot for the significant *fmn* ZT 12 vs All model. Classes were defined as ZT 12 and All (ZT 0, ZT 4, ZT 8, ZT 16, and ZT 20). **e,** Corresponding loadings plot for the *fmn* ZT 12 vs All OPLS-DA model. Isotopologues are represented by shapes, metabolites by color and size of the shapes represents significance as defined through VIP values. Abbreviations as defined in Fig. 3g. **f,** SUS plot comparing the loading plots from the *fmn* ZT 0 vs All (**c**) and the *fmn* ZT 12 vs All OPLS-DA models (**e**) with a regression line and equation. Green refers to VIP compounds in both models, orange refers to VIP compounds only in the *fmn* ZT 0 vs All model, and blue refers to VIP compounds only in the *fmn* ZT12 vs All model. Shapes refer to isotopologues. **g,** SUS plot comparing the loadings plot from the *fmn* ZT 0 vs All (c) and the WT ZT 4 vs All (Fig. 3g) OPLS-DA models with a regression line and equation. Green refers to VIP compounds in both models; blue refers to VIP compounds in the WT model; red refers to VIP compounds in the *fmn* model. Shapes refer to isotopologues. **h,** A simplified overview of downstream glucose metabolism indicating potential routes for the biosynthesis of VIP metabolites highlighted in f according to time point. Isotopologue text color refers to time point.

### Only the dawn biosynthetic ‘rush hour’ is similar to WT ‘rush hour’

Given that *fmn* had two peaks with strong biosynthetic activity, we then investigated whether excess glucose metabolism at ZT 0 and ZT 12 followed a similar set of pathways in *fmn* using a shared and unique structures (SUS) plot (Fig. 5f). In this analysis, the loadings (or distribution of isotopologue enrichments) from both models were plotted against each other with ZT 0 vs All (Fig. 5c) on the y-axis and ZT 12 vs All (Fig. 5e) on the x-axis. The resulting plot highlighted a weak correlation between the two time points through both unique (blue and orange), as well as conserved (green) metabolism (Fig. 5f). For example, enrichments of ornithine M+1 and alpha-ketoglutarate M+2 were relatively higher at both ZT 0 and ZT 12 but were only significantly contributing to the separation of ZT 12 from other time points. On the other hand, enrichments of isotopologues such as glutamate M+2 and M+4 only significantly contributed to the separation of the ZT 0 time point. From both models, broad labelling of downstream glucose products with differences in metabolite and/or labelling were noted at both time points. Overall increased TCA cycle labelling was apparent at both time points through unique and common isotopologues while glucose carbons may be incorporated into glycolysis more at ZT 12 (Fig. 5h). Overall, this highlights that although a few similar pathways may be activated in *fmn* at the two LD transitions (or dawn and dusk), generally different isotopologues are generated (i.e. unique metabolic pathways used).

Since *fumin* at ZT 0 and ZT 12 and wild type at beginning of the light phase (ZT 4) had all shown increased biosynthesis from labelled glucose, we next looked to determine whether *fmn* used glucose similarly to WT at these time points. When comparing the loading plots from the early light period from both models (eg WT ZT 4 vs All (Fig. 3g) and the *fmn* ZT 0 vs All (Fig. 5c), the resulting SUS model highlighted a notable degree of correlation between the two conditions (Fig. 5g). Furthermore, several conserved compounds significantly contributed to the separation of both ZT 4 in WT and ZT 0 in *fmn*, highlighted in green. For example, proline M+4 and glutamine M+3, M+4, and M+5 enrichments were relatively higher at both ZT 4 in WT and ZT 0 in *fmn* relative to other time points. On the other hand, compounds significantly contributing to the separation of *fmn* ZT 0 (VIP compounds such as citrate M+3, M+5, and M+6) were also noted and although they displayed similar changes in WT, they did not meet the significance cutoff as defined by a VIP value greater than one. In contrast, fewer concerted changes were observed between *fmn* at ZT 12 and WT at ZT 4 (Extended Data Fig. 4c). This analysis suggests that there is a similar and specific set of metabolic pathways activated at ZT 4 in WT and ZT 0 in *fmn* compared to other timepoints indicating a potential phase advancement in *fmn* of metabolic pathways primed to utilize glucose in the early morning period.

### Increased diurnal and ultradian biosynthetic rhythms in fumin

Having observed unique processing of glucose at two time points in *fumin*, we next asked if the changes are due to underlying cycling over the course of a day. To address isotopologue rhythmicity, 20-28-, 24-, and 12-hr periods were tested. Overall, a greater number of rhythmic isotopologue enrichments and/or pools were observed in *fmn* than WT for all periods tested across different RAIN and JTK FDR cut-offs (Fig. 6a, Extended Data Fig. 5a, and Fig. 6e). For further analyses, compounds meeting a 0.2 RAIN FDR cut-off were used (Fig. 6b, Extended Data Fig. 5b, and Fig. 5f). In contrast to WT, metabolite pools cycled with 20-28-, 24- and 12-hr periods. Most of the significant 24-hr cycling labelled isotopologue enrichments were noted to peak around ZT 0 to ZT 4 while unlabeled forms peaked during the dark period between ZT 16 and ZT 20 and pools were noted to peak around ZT 4 (Fig. 6c). Since the peak in overall metabolite levels (pools) and enrichments (labelled isotopologues) both occurred between ZT 0 and ZT 4, this suggested that these arose from the incorporation of labelled glucose. However, alternative sources of carbon contribution cannot be ruled out. This trend was also observed in phases of significant 20-28-hr cyclers (Extended Data Fig. 5c). Similar to the observed 24-hr rhythmicity, significant compounds with 12-hr periods also showed a number of labelled isotopologues with peaks between ZT 0 and ZT 4 along with pools peaking at ZT 4 and non-labelled forms, M+0, peaking around ZT 8 (Fig. 6g). In this case, because pools are peaking in between labelled and unlabeled forms, it is difficult to determine whether overall pool levels are increased due to biosynthesis from glucose. However, it is likely that glucose carbons contributed to the increased pool levels noted as alternative routes are limited. Overall, rhythmicity analyses support increased biosynthesis of isotopologues at the LD transitions in *fmn* with a substantial contribution to the increased biomass arising from glucose carbons. Furthermore, the peak of rhythmicity around ZT 0 in *fmn* also supports the idea of phase advanced biosynthesis from WT.

**Fig. 6:**
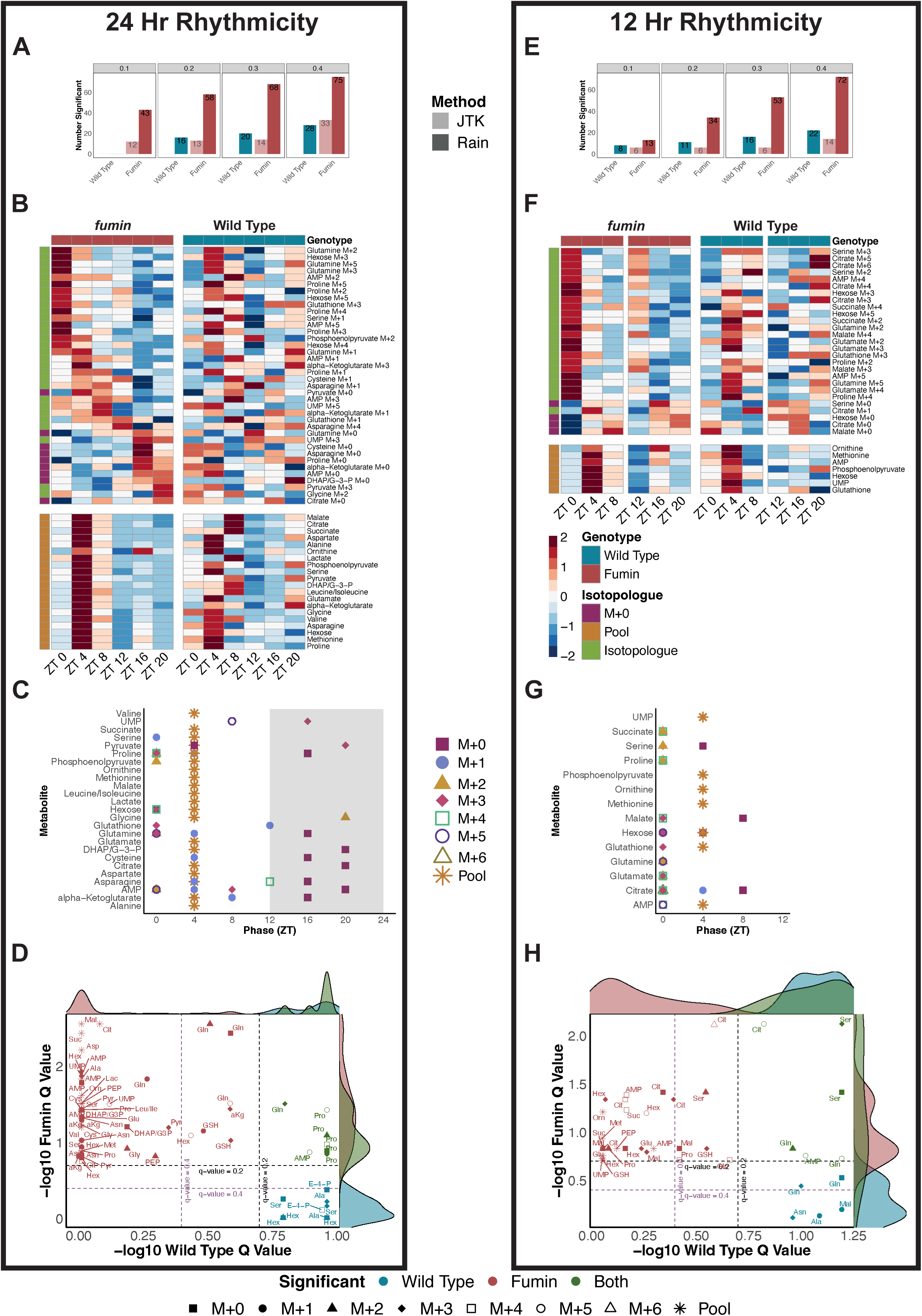
Increased diurnal and ultradian biosynthetic rhythms in *fumin.* **a,e,** Overview of the number of significant 24-hr (a) and 12-hr (e) rhythmic compounds observed at RAIN (dark) or JTK (light) q-value cut-offs of 0.1, 0.2, 0.3, and 0.4. **b,f,** Phase ordered heatmaps of significantly cycling compounds with periods of 24-hr (b) or 12-hr (f) in *fmn* at a RAIN q-value cutoff of 0.2. Enrichments for WT are shown for comparison. Legend for both heatmaps is shown with f. **c,g,** Distribution of RAIN phases for significant 24-hr (c) and 12-hr (g) compounds in *fmn* grouped by metabolite. Shapes and colors refer to isotopologues or pools. **d,h,** Comparison of 24-hr (d) or 12-hr (h) significant cycling compounds in WT and *fmn* as defined by a RAIN q-value less than 0.2. q-value cutoffs of 0.2 (black dotted line) and 0.4 (purple dotted line) are shown. Color refers to significant rhythmicity in one or both genotypes while shapes refer to isotopologues.

Since rhythmicity was observed in both genotypes, we next explored whether similar compounds were cycling in WT and *fmn*. For 24-hr rhythmic compounds, about 50% of the isotopologues cycling in WT were also noted to be 24-hr rhythmic in *fmn* (Fig. 6d). A similar observation was also noted with 20-28 hr rhythmic compounds (Extended Data Fig. 5d). In contrast, *fmn* primarily contained unique isotopologues and/or pools that were rhythmic, but it is important to note that a few are trending towards significance in WT (glutathione M+1 and M+3 and glutamine M+0 and M+2). A similar trend was also noted for 12-hr rhythmic compounds (Fig. 6h). However, glutamine M+0 and M+3 were significant in WT for 12-hr periods but trended towards significance in *fmn*. This highlights that a majority of the cycling observed in *fmn* was uniquely gained rather than conserved or lost from the cycling observed in WT. Overall, rhythmicity analyses also supported TCA cycle labelling in *fumin* with contributions from both diurnal and ultradian rhythmicity while cycling of glycolysis and PPP related isotopologues was primarily diurnal.

### Shift from ultradian to diurnal rhythmicity is observed in product to precursor relationships in fumin compared to wild type

In order to understand how carbon flow was occurring through the TCA cycle, we employed a product to precursor relationship analysis^40^. This analysis generates ratios (phi values) using different mathematical relationships between TCA cycle precursors and products to provide further insight into how glucose is being used. For example, the SM4 ratio looks at the formation of malate M+4 from succinate M+4. Since malate M+4 is only derived from succinate M+4, the ratio provides insight into step-wise flow of oxidative carbons at this TCA cycle step relative to other pathways like anaplerotic pyruvate carboxylase. Neither of the inputs for this ratio displayed significant rhythmicity (24- or 12-hr), however, a 12-hr rhythm trending towards significance was observed in the WT ratio (Fig. 7a). This highlights the additional value gained from the ratio analysis which was not captured at the enrichment-level alone and suggests rhythmicity between oxidative and synthetic mitochondrial metabolism.

**Fig. 7:**
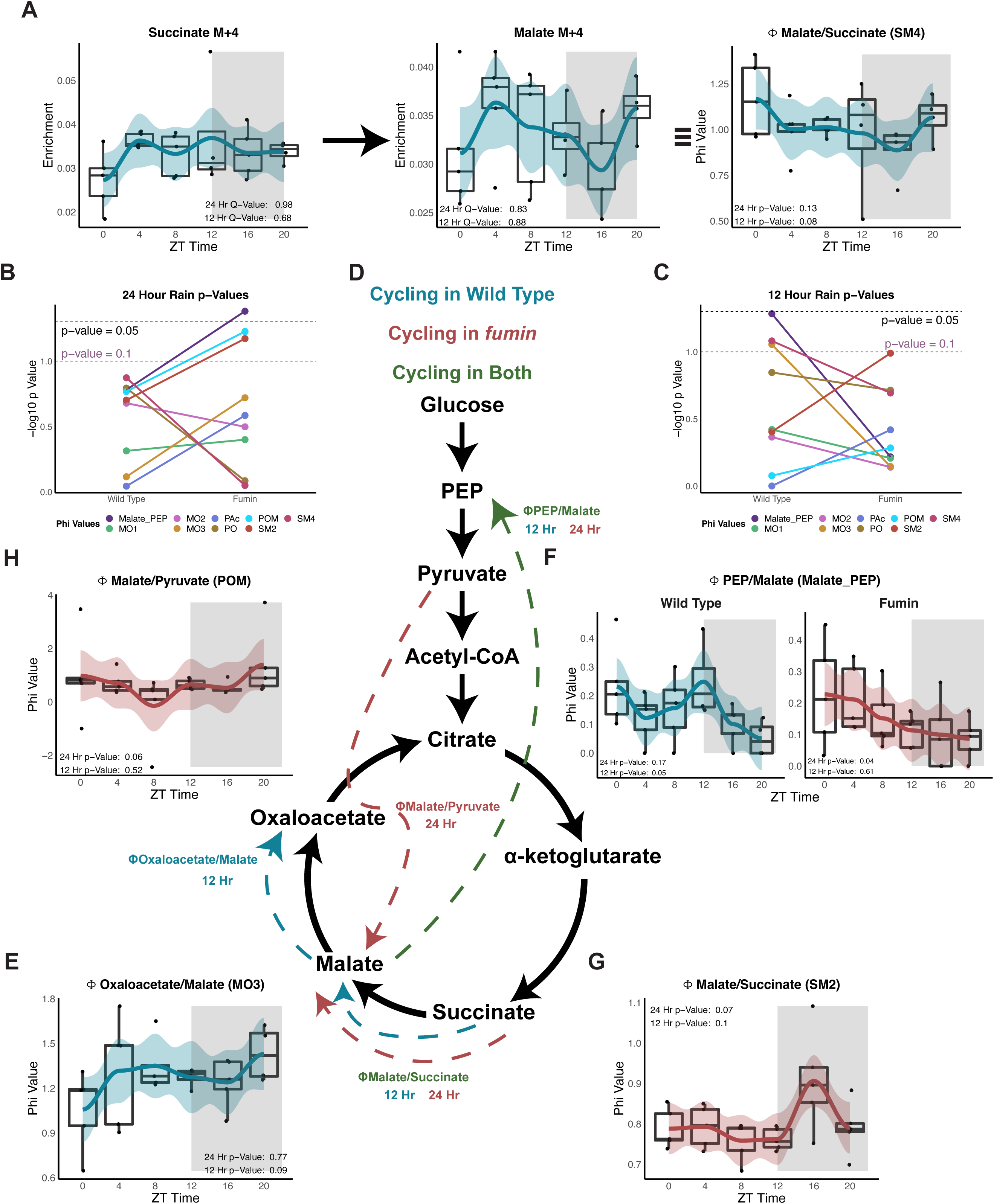
Shift from ultradian to diurnal rhythmicity is observed in product to precursor relationships in *fumin* compared to wild type. **a,** Box plots with LOESS traces for a representative product to precursor relationship (malate M+4/succinate M+4) in WT (SM4, right panel), the precursor, succinate M+4 (left), and the product, malate M+4 (middle). N = 4 to 5 biological replicates of 15 male flies per time point. Rhythmicity was tested for 24-hr and 12-hr periods using RAIN and results with a p-value less than 0.1 are shown. **b,c,** Summary of RAIN p-values for 24-hr (b) and 12-hr (c) periods for WT and *fmn*. Colors represent product to precursor relationships (phi values). A p-value cut-off 0.05 is shown in the black dotted line a p-value cut-off of 0.1 in the purple dotted line. **d,** Overview of significant phi value relationships. **e-h,** Box plots and LOESS traces for phi values with p-value less than 0.1 in WT and/or *fmn*. n = 4 to 5 biological replicates with 15 male flies per time point

Overall, the Malate_PEP phi value demonstrated significant 24-hr rhythmicity and two others, POM and SM2, trended towards significance in *fmn* (Fig. 7b and d). In contrast to 24-hr rhythms, 12-hr cycling trended towards significance in three WT phi values (SM4, MO3, and Malate_PEP) (Fig. 7c and d). For SM4, a ratio of one (i.e., ZT 4) indicated the formation of malate M+4 from succinate M+4 was not diluted by non-oxidative carbon flows. However, a ratio greater than one (i.e., ZT 20) could indicate either reversal of the pathway or the entry of alternately labelled carbons in the formation of malate (Fig. 7a middle panel). Similarly, ratios greater than one in the 12-hr rhythmic MO3 phi value suggested that either the malate was being formed from oxaloacetate or alternative routes with labelled carbons were being used to form oxaloacetate (Fig. 7e). Given that fly bodies (and not single tissues) were examined and that some compartmentation is to be expected, then phis become more semiquantitative or qualitative indices of flux since true single pool steady state assumptions may not apply. Interestingly, the Malate_PEP ratio switched from 12-hr rhythmicity in WT to 24-hr rhythmicity in *fmn* (Fig. 7f) and indicates a relative shift in glycolysis vs gluconeogenesis. However, the ratio was less than one in both genotypes indicating that unlabeled carbons from alternate metabolites, such as glucose, were diluting the formation of PEP from malate. A different ratio looking at the formation of malate from succinate using the M+2 forms, SM2, displayed 24-hr rhythmicity in *fmn* with values less than one indicating the entrance of alternate sources of unlabeled carbons (Fig. 7g). *fmn* also displayed 24-hr rhythmicity in the POM ratio which looked at the formation of malate from pyruvate (Fig. 7h). Here, in instances where the ratio decreased less than 1, it implied alternative sources of unlabeled carbons diluting the pathway. Overall, the product to precursor analysis highlighted a shift in key oxidative and synthetic metabolic flows from ultradian rhythmicity in WT to 24-hr rhythmicity in *fmn* pointing not only to differences in glucose utilization but also potential differences in demand.

### Feeding rhythms and a 4-hour short-term fast do not impact processing of excess glucose

Since timing of food availability can be a zeitgeber for molecular clock rhythmicity, we asked how nutrient factors impact the time shift in biosynthesis observed between WT and *fmn*. To determine whether altered feeding rhythms between the genotypes were impacting downstream glucose biosynthesis, feeding profiles of WT and *fmn* were monitored through the ARC assay revealing similar levels of food consumption in both genotypes overall and during the light and dark periods. However, the feeding peak in *fmn* (ZT 4.5) was phase delayed in comparison to WT (ZT 3.25) (Fig. 8a). Since the biosynthesis peak in *fmn* was observed to be phase advanced from WT while the feeding rhythms were phase delayed, we concluded that overall feeding was not an underlying cause of the biosynthesis shift in *fmn*.

**Fig. 8:**
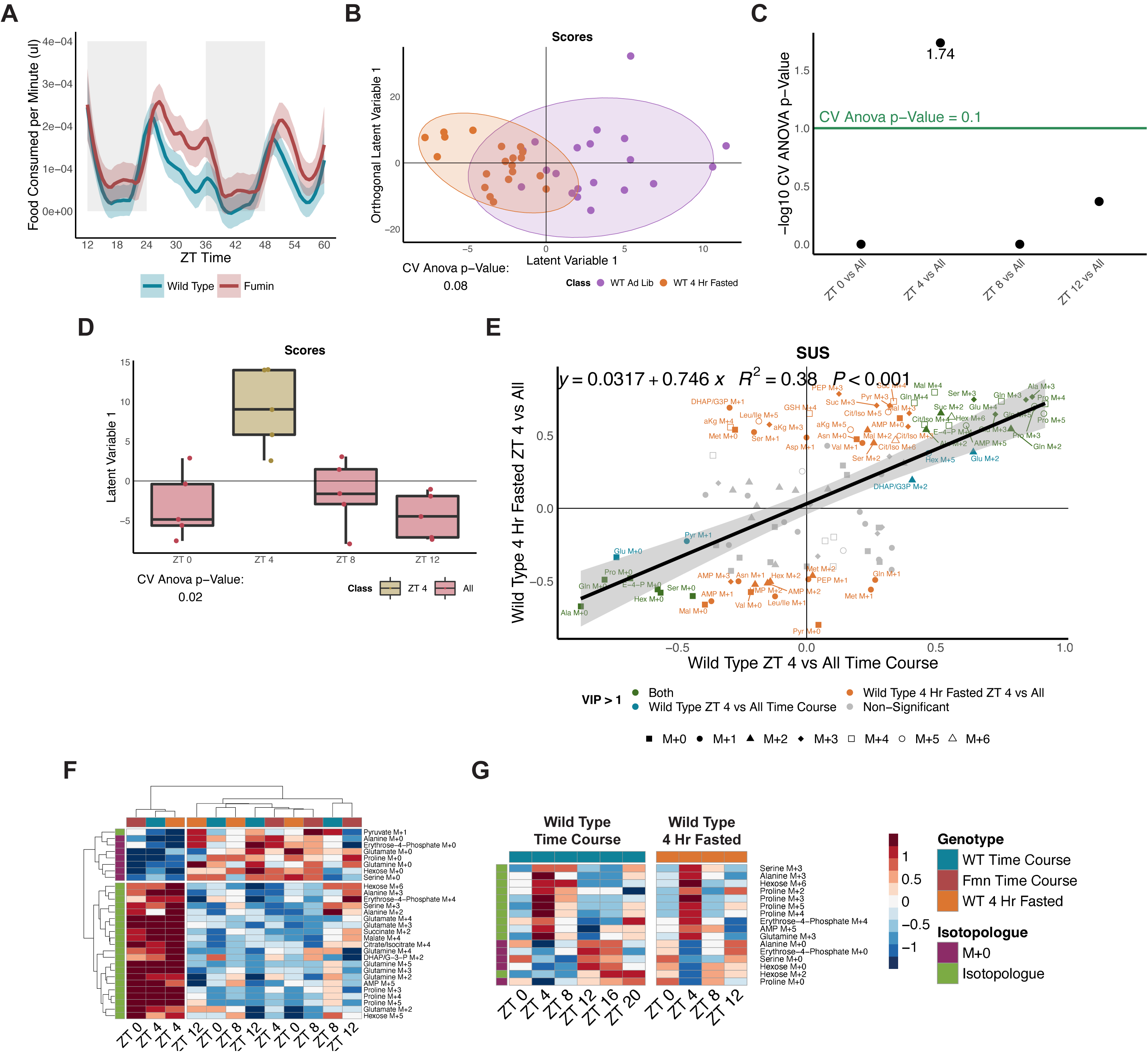
Feeding rhythms and a 4-hour short-term fast do not impact processing of excess glucose. **a,** LOESS trace of feeding rhythms measured by volume of food consumed per minute by WT and *fmn* flies across 2 days using the ARC assay. n = 53 to 62 per genotype. **b,** A two-component OPLS-DA model comparing WT ad libitum and short-term fasted flies. All time points collected (ZT 0, 4, 8 and 12) are included for each condition. N = 5 biological replicates with 18 to 24 flies per time point and group. **c,** Negative log CV-ANOVA p-values for each time point tested in pairwise OPLS-DA models in the short-term fasted condition. Significance was defined by as a CV-ANOVA p-value less than 0.1 (shown as points above the green line). **d,** Scores plot for the significant short-term fasted ZT 4 vs All OPLS-DA model. Two classes were defined as ZT 4 and All (ZT 0, 8, and 12). **e,** SUS plot comparing the loadings from the WT ZT 4 vs All (Fig. 3g) and the short-term fasted ZT 4 vs All (Extended Data Fig. 6b) OPLS-DA models with a regression line and equation. Green refers to VIP compounds in both models; orange refers to VIP compounds in the short-term fasted model; blue refers to VIP compounds in the WT model (Fig. 3g). Shapes represent isotopologues. **f,** Clustered heatmap on both the x and y-axes shown using ZT 0 to 12 time points from earlier WT and *fmn* samples along with the short-term fasted samples for VIP compounds determined from the WT ZT 4 vs All model (Fig. 3f and g). **g,** Phase ordered heatmap showing levels of 24-hr rhythmic metabolites from wild type (Fig. 3b) for the short-term restricted time course.

To further establish whether the availability of food was driving biosynthesis in WT, a shortened time course from ZT 0 to ZT 12 was analyzed under *ad libitum* and a 4-hour fasted condition, where food was withheld for 4 hours prior to injection with the glucose tracer. Distinct metabolic profiles were noted for the two conditions, with different labelled isotopologue enrichments associated with each (Fig. 8b and Extended Data Fig. 6a). We next determined whether ZT 4 was still metabolically distinct from remaining time points under the short-term fasted condition. The ZT 4 model did emerge as significant in a pairwise discriminant model to all remaining time points (Fig. 8c and d). Similar to the earlier observation, a majority of the labelled isotopologues were associated with ZT 4 (Extended Data Fig. 6b) pointing to increased biosynthesis from glucose. To further determine whether the presence of increased biosynthesis was arising from similar compounds and/or pathways observed previously (Fig. 3g), we performed a SUS analysis using the loadings from the both ZT 4 vs All (WT -Fig. 3g and short-term fasted – Extended Data Fig. 6b) models. The resulting plot showed conserved and similar changes occurring at both ZT 4 time points (highlighted in green) (Fig. 8e). It is important to note that several compounds uniquely contributed to the separation of ZT 4 in the short-term fasted model, but this is to be expected as food restriction would impact metabolism and excess glucose processing. We also performed a complementary analysis using the enrichments identified from the WT ZT 4 vs All model (Fig. 3g) as significantly contributing to the separation of ZT 4 from all other time points (VIP greater than 1). Using these VIP metabolites in a clustering analysis with calculated enrichments from ZT 0 to 12 from the WT, short-term fasted WT and *fmn* time courses, two main clusters emerged for both time and compounds (Fig. 8f and Extended Data Fig. 6c). On the time axis, ZT 4 from both the WT and short-term fasted condition clustered with the *fmn* ZT0 time point while all remaining time points formed a second cluster. On the metabolite axis, labelled and unlabeled isotopologues formed the two main clusters. This clustering further highlighted the peak in labelled isotopologue enrichments at ZT 4 and ZT 0 in WT and *fmn,* respectively. Since rhythmicity analyses had also previously shown peaks in labelled isotopologues and troughs for unlabeled forms at ZT 4, we assessed the overall enrichment profiles of the short-term fasted condition using the 24-hr rhythmic metabolites (identified in Fig. 3b). This further showed similar patterns across the two data sets (Fig. 8g). Overall, this highlighted sustained increased downstream biosynthesis from glucose carbons and peaks in rhythmicity at ZT 4 in WT irrespective of food pointing to a potential underlying circadian driver rather than the availability of food.

## Discussion

Glucose and glucose metabolism play a central role in energy metabolism. Glucose levels are closely regulated and maintained and disruptions to glucose metabolism lead to diseases such as diabetes^41,42^. Circadian rhythms have not only been observed in glucose levels but also in processes involved in nutrient metabolism^43^. Moreover, glucose homeostasis has been shown to be impacted by deletion of a core clock component, *Bmal1*, both at a global and tissue-specific level in mice. For example, global deletion of *Bmal1* has been associated with impaired glucose tolerances and gluconeogenesis while tissue-specific deletion of *Bmal1* in the liver and intestine resulted in hypoglycemia and impaired glucose absorption, respectively^44–47^. This work demonstrated overall time of day differences and the presence of rhythmicity in glucose related metabolites in blood samples from human subjects. Based on the human data, we used glucose as a tracer to begin exploring time-of-day differences in how time-dependent metabolic programming occurs in WT in comparison to the *fumin* hyperactive mutant. Although trehalose is a major carbohydrate in *Drosophila,* glucose is regulated by physiological processes and environmental cues^48^. Additionally, *Drosophila* contain homologs to mammalian insulin, Drosophila insulin-like peptides, insulin receptor, and insulin substrate. Because carbohydrate metabolism and signaling pathways are conserved across humans and *Drosophila, Drosophila* serves as a reasonable model to understand how glucose is processed at different times of day^49,50^.

By using labelled glucose as a tracer, we have detected time-of-day differences in the fate of glucose carbons *in vivo* in *Drosophila.* The method used here to introduce a bolus of glucose and sample after allowing an hour for metabolism to occur may not achieve complete steady-state labelling of metabolites downstream of glucose, thus a pseudo-steady state condition is achieved which allows comparisons across times and groups to be performed. We previously reported significant 20-28-hr total metabolite pool rhythmicity in wild type under LD cycles of asparagine, glutamate, citrate/isocitrate and fructose^25^; these pools and their isotopologues were not significantly rhythmic in this study. Furthermore, cycling was not observed in metabolite pools for the isotopologues that were significant in WT. We rationalized that this difference is the result of the glucose challenge in the current study. A number of glucose-related pathways (TCA cycle, PPP, and glycolysis) were noted to use labelled glucose as indicated by the higher relative labelled isotopologue enrichments and peaks of cycling isotopologues at ZT 4. Furthermore, recent work has shown that mass action relationships primarily dictate the regulation of metabolites^51^. This supports the results observed in WT since different pathways of processing glucose are activated. However, we also observe that there is a time-of-day dependent effect where these pathways may be more active at ZT 4 allowing glucose to be cleared in a potentially more efficient manner than at other times. As glucose is sensitive to environmental cues^48^, this therefore implies that underlying processes for glucose metabolism are primed at ZT 4 leading to the rhythmicity and synthesis of specific isotopologues which is not reflected in metabolite pool levels. Therefore, although the introduction of a glucose bolus impacts metabolism, the activated pathways may still be used in steady state studies and mask rhythmicity at the pool-level. This highlights the importance of understanding how specific metabolite isotopologues are altered, which can have significant implications for understanding the interaction of the circadian clock and metabolic disease.

On the other hand, the number of rhythmic compounds in *fmn* was increased when compared to WT and included both isotopologues and pools. It is important to note that the presence of cycling in metabolite pools can convolute the interpretation of whether increased biosynthesis of rhythmic isotopologues is occurring specifically for the isotopologue of interest or generally of the pool. However, for 24-hr rhythmic compounds, the peak in metabolite pools aligned more closely with the peak observed in isotopologues and thus could point to the production of the isotopologues driving the rhythmicity of the pools rather than the synthesis of the metabolite through alternate pathways (other than from glucose) resulting in unlabeled forms. Further studies would be needed to definitively discriminate whether labelled carbons from metabolites other than glucose were affecting the pool sizes.

The increase in TCA cycle labelling in *fmn* may be a result of the increased activity, an increased metabolic rate^52^, a switch in fuel preference and/or decreased sleep observed in *fmn* which may necessitate an increased need for TCA intermediates and/or amino acid production. Moreover, the lack of PPP labelling in *fmn* in relation to WT overall may reflect an absence or inability to properly regulate redox oscillations as this pathway produces NADPH^53^ or could simply reflect decreased glucose phosphorylation, increased gluconeogenesis and/or increased glycogenolysis. Dysfunction in the PPP can have implications for the behavior and rhythms in *fmn*. Furthermore, *fmn* was noted to process glucose differently at the LD transitions through the pathways used as well as the isotopologues generated. Similar to elevated TCA cycle labelling observed in *fmn* irrespective of time when compared to WT, glycolytic and TCA cycle related metabolites were also observed to be labelled at both ZT 0 and ZT 12 although a number of unique isotopologues to both time points were also noted. In addition to concentration being a driving factor of consumption flux^51^, our results point to differences in glucose consumption pathways dependent on time-of-day and/or genotypes. Furthermore, increased energetic demands arising from hyperactivity in *fmn* may also be driving not only increased glucose utilization but also differences in glucose processing by time of day.

The striking robustness of the metabolic biosynthesis ‘rush hours’ via isotopologue enrichments observed at ZT 4 in WT (ad lib or fasted) and ZT 0 in *fmn implies* that while *fmn* is slightly phase advanced in its morning glucose catabolic ‘program’, the overall program remains the same. Because *Drosophila* consume food primarily during the light period^54^, one possibility for the phase advancement in catabolism is altered underlying feeding rhythms. Mistimed feeding can serve as a potent zeitgeber especially for peripheral clocks with impacts on metabolic processes^55^. We observed however, the opposite result as feeding rhythms in *fmn* were phase delayed compared to WT while the biosynthesis peak was phase advanced, feeding therefore was not the primary driver of the isotopologue rhythmicity observed. This also begins to deconvolute the influence of feeding on downstream glucose catabolism from that being driven by the presence of excess glucose and/or primed pathways. As feeding acts as a potent zeitgeber especially in peripheral clocks, misaligned feeding which can occur in shift work has been associated with altered metabolism and adverse health consequences^55,56^, however; the observed phase advancement of biosynthesis also highlights the importance of understanding whether shifts in metabolism become decoupled from feeding and understanding the implications that may arise from this.

Product to precursor relationship analysis provides further insight into how glucose was being used in a manner that was not captured at the isotopologue level. For example, the presence of rhythmicity in a ratio was not necessarily preceded by cycling in the individual inputs. Through this analysis, we were able to observe an overall shift from 12-hr rhythmicity in WT to 24-hr rhythmicity in *fmn* which followed the activity patterns of the genotypes. Specifically, in WT, we were able to follow the flow of carbons in a step-wise manner from succinate M+4 to malate M+4 to oxaloacetate M+4 through two ratios, SM4 and MO3. Both ratios looked at oxidative reactions within the TCA cycle and were around one indicating step-wise carbon flow through the reactions. However, at ZT 0 and ZT 20, the SM4 ratio was higher than one implying compartmentalization across different tissues, some error in the measurement, or incorporation of labeled bicarbonate during pyruvate carboxylation. On the other hand, *fmn* also displayed rhythmicity in oxidative reactions through the phi Value SM2. This value used M+2 forms of malate and succinate and implied that the pathway was being diluted by alternately labelled or unlabeled carbons except at ZT 16 where the reaction was closer to a direct relationship between the product and precursor. Additionally, the POM phi value was an indicator of anaplerosis driven by pyruvate carboxylase. This phi value was close to one at ZT 0 and ZT 20 in *fmn* implying that oxaloacetate was primarily being formed through anaplerotic pathways while at the other time points, alternate sources such as malate and/or the introduction of aspartate could be contributing to the generation of oxaloacetate. Taken together, the phi value product to precursor relationship analysis provides additional insight into how glucose is being used in both genotypes that is not gained from typical analysis using enrichments.

While the isotope tracing portion of this study was performed in Drosophila, we speculate that there are important implications for understanding glucose and energy response in humans. For example, increasing evidence suggests that caloric intake biased towards earlier times have health benefits with regards to weight loss, glycemic control^57,58^, and cardio-metabolic outcomes^59^. A significant challenge in the clinical application of this knowledge is overcoming lifestyle and habitual resistance changing or limiting time windows for food intake. For example, dinner is an important social dinner component, while skipping breakfast is generally not recommended creating a difficulty in creating realistic food timing windows. Similarly, with regards to activity, overweight individuals under an acute high fat diet have been shown to benefit from exercise with respect to cardiorespiratory fitness irrespective of timing, but only glycemic control is improved by evening exercise^60^. By understanding the downstream mechanisms at work through studies such as the current work, we may be able to eventually expand the temporal window of effectiveness of food or exercise through dietary or pharmacology intervention to more effectively employ these strategies.

Overall, we have shown the sensitivity of the metabolic fate platform to detect low-level differences in downstream glucose metabolites at different times of the day. This approach can be adapted for higher time resolution studies and/or other labelled tracers in *Drosophila* but also in other model organisms and potentially humans due to the small amount of tracer required and high mass sensitivity of the platform. In our current study design, up to one hour resolution can be achieved as this is the time needed for glucose to be sufficiently incorporated into downstream metabolites. More generally, this approach can be used to explore how circadian regulation of metabolism occurs in conjunction with other questions related to disease pathophysiologies, sleep, and influence of other environmental zeitgebers. Using the resulting insights, a better understanding of metabolic changes can be gained which can drive interventions in a more precise manner both in terms of targets as well as timing.

## Acknowledgements

We thank Lisa Bottalico for discussions on LCMS methodology; Gregory Grant for discussions on considerations for proper sample randomization; Tom Brooks and Antonijo Mrcela for discussions on cycling measurement algorithms; Sunil Dhakad, Maheswari Karthikeyan, Shefali Lathwal, and Raghav Sehgal (Elucidata) for their discussions and help on performing natural abundance corrections and phi value calculations; Thomas Jongens for suggestions and editing; and Joeseph Bedont for discussions around short sleeping mutants. This work was supported by NIDDK of the National Institutes of Health under award number R01-DK120757 to A.S. and A.M.W. D.M.M. and S.D.R. were supported through a Pharmacology T32 Training Grant (T32 GM008076). A.S. is an investigator of the Howard Hughes medical Institute (HHMI).

## Author Contributions

Conceptualization, D.M.M., S.D.R., A.S., and A.M.W.; Methodology, D.M.M, S.D.R., A.B.; Formal Analysis D.M.M., S.D.R., and A.M.W.; Investigation, D.M.M, S.D.R., S.L.Z., and P.H.; Resources, S.L.Z., A.B., P.H., R.G.K., and A.S.; Writing – Original Draft, D.M.M., S.D.R., and A.M.W.; Writing – Review & Editing, D.M.M., S.D.R., S.L.Z., A.B., P.H., R.G.K., P.K., A.S., A.M.W; Visualization, D.M.M, S.D.R., A.M.W.; Supervision A.M.W.; Funding Acquisition, A.S. and A.M.W.

## Methods

### Drosophila Strains

Drosophila melanogaster, wild type (isogenic w^118^ stock) and *fumin* mutants as characterized previously^37^, were maintained on standard cornmeal/molasses medium at 25°C under 12:12 LD conditions.

### Geotaxis and Locomotion Assays

After three days under LD entrainment, 5-to-7-day-old wild type or *fumin* flies were anesthetized on ice for 15 minutes at zeitgeber time (ZT) 0 (lights on) or 12 (lights off). Flies were either not injected (control) or injected with PBS or 1M Glucose. For geotaxis assays^61,62^, flies were placed in new vials without food for one hour after which climbing activity was measured by the time taken to climb 4 cm per fly, measured in triplicate using 5-8 flies per group. For locomotion assays^63,64^, flies were placed in 5 x 65 mm glass tubes with 5% sucrose post-injections and monitored using Activity Monitoring System devices (DAMS) from Trikinetics (Waltham, MA) for a minimum of 24 hours in 25°C LD incubators. Locomotion data was analyzed using a custom MATLAB (MathWorks, Natick, MA Versions 2013B and 2020A) script^65^ with additional analysis and plotting performed in R (version 4.0.3). Comparisons for both assays were made using a two-sided t-test using p<0.05 as a threshold for significance.

### Feeding Assay

An Activity Recording CAFE (ARC) assay^66^ was used to record feeding rhythms of wild type and *fumin*. Male flies were entrained under 12 hr Light:Dark conditions within one day of eclosion at 21°C. On day 6, entrained flies were used to setup the feeding assay with one fly per well containing 300 µl of 2% agar. Genotypes were loaded in alternating wells. A capillary containing a (infrared) dye (for detection of food level) and 2.5% yeast and 2.5% sucrose was placed into each well as a food source. After allowing for habituation to the setup, data was recorded for 48 hours with capillaries replaced each day before ZT 12. This assay was performed at 21°C to minimize evaporation of the capillaries. Data was visualized in R (version 4.0.3) and any wells containing flies that had not survived, evaporation only of food and/or inconsistent food level recordings were discarded. Data was processed using a python program (Noah15.2) with additional analyses in R (version 4.0.3).

### Glucose Injection Time Courses

Within one day after eclosion, 15 to 18 males were sorted into a new vial for entrainment to LD rhythms for a minimum for 3 days in appropriate incubators at 25°C. All flies were between 5-7 days old at the time of injection. One vial was placed on ice to immobilize flies 15 minutes prior to each injection time point. The injection solution consisted of 1M uniformly labeled (^13^C_6_) glucose (CIL D-Glucose, 99%) and a blue dye (50x dilution of McCormick, FD&C Blue Dye No. 1) to allow for visualization during injections and normalization for variations in injection volumes during data processing. The solution was injected into the thorax using glass capillaries (9 cm x 1.14mm diameter, Drummond Scientific, Broomall, PA) on ice. After injection, flies were moved to empty vials containing a kimwipe moistened with 1 ml of water to prevent flies from becoming dehydrated and returned to the appropriate light or dark 25°C incubator to metabolize the glucose tracer. After one hour, flies were collected on dry ice and stored at -80°C until extractions. Five separate days of injections were performed as biological replicates for both WT and *fumin* mutants. Any flies which died after the injections were discarded. No difference in survival rate (>95%) was noted between the genotypes.

For the shortened ad libitum versus 4-hour short term fasted time course, injections were modified as follows: 20 to 24 male wild type flies were collected and entrained within one day of eclosion and were injected on day 7. Four hours prior to each time point, flies were transferred either to fresh food vials (ad libitum condition) or agar vials (4-hour fasted condition). Each group was injected with either 1 M ^13^C_6_-glucose or 1 M ^12^C_6_-glucose (Acros Organics alpha-D(+) Glucose, 99%) with blue dye (504 µM Sigma Aldrich Brilliant Blue FCF, analytical standard) dissolved in PBS.

### Metabolite Extraction and LC-MS Measurements

Fly heads and bodies were separated prior to metabolite extraction, using an adaptation of the Bligh-dyer extraction^38,67^. Briefly, to each fly body sample, a total of 600 µL of cold 2:1 methanol:chloroform was added and homogenized in a bead-based tissue homogenizer at 25 Hz for 4 minutes (TissuLyser II, Qiagen, Hilden, Germany). 200 µL each of water and chloroform were then added, followed by centrifugation at 18787xg for 7 to 10 minutes at 4°C. 350 to 400 µL of the upper (aqueous) layer was collected and dried under vacuum until dry or overnight. Samples were resuspended at 4 µl/fly body for negative mode and diluted to 5.33 µl/fly body for positive mode using 50:50 water:acetonitrile. Each sample was analyzed separately in ESI positive and negative modes. Transitions included in each method are shown in Table S1 and S2. Chromatographic separations utilized an ACQUITY UPLC BEH Amide column (2.1 x 150mm, 1.7 µm) on a Waters ACQUITY H-Class UPLC coupled to a triple quadrupole Waters Xevo TQ-S Micro MS (Milford, MA). LC conditions for the positive-ionizing method were performed as described previously^68^ with a modified gradient from 100 to 20.6 % B over 15 min at 0.35 mL/min, followed by a wash of 100% A for 5 min. Mobile phase B was changed from 0 to 100% from 20 to 22 min and held for column equilibration until 30 min. For the negative-ionizing method, solvents consisted of 95:5 water:acetonitrile with 20mM ammonium bicarbonate, pH 9 (mobile phase A) and 90:10 water:acetonitrile with 20mM ammonium bicarbonate, pH 9 (mobile phase A). The gradient was changed from 100 to 20.6% B over 15 min at 0.4 mL/min, followed by a wash of 100% A for 5 min. Mobile phase B was changed from 0 to 100% from 20 to 22 min and held for column equilibration until 30 min. Ion counts were acquired through multiple reaction monitoring (MRM).

For the feeding time course samples, the LCMS analyses was performed as described above with the addition of 5 µM ammonium phosphate to both mobile phases in both ionization modes.

### Data Processing and Normalization

Chromatograms were processed using TargetLynx under MassLynx version 4.1 or using El-MAVEN (version 0.12.1 beta) and exported as ion counts for further processing in R (version 4.0.3). Column pressure deviations were monitored to identify poor sample injections which were excluded from further analysis. MRMs which were either not detected or overlapped with other chromatographic peaks and thus prevented unequivocal peak integration were set to zero. For each sample, analytical duplicates were injected in a randomized order, along with QC samples every 4 to 10 injections, which was comprised of a pool of all samples.

In order to perform an instrumental drift correction, consecutive QCs were averaged. Next, ion counts were used to calculate metabolite pools and were also used to calculate a ratio of each transition acquired to the calculated pool. A LOESS correction was performed on the metabolite pools to minimize the impact of differences in detection of isotopologues across samples^69^. The corrected pools were multiplied by the ratios calculated earlier to regenerate transition-level data corrected for instrumental drift. A validated FCF transition from the negative mode data was used to normalize for variation in glucose injection volumes for the wild type and *fumin* time courses. Analytical replicates were then averaged and used to perform a natural abundance correction using the Polly Phi LCMS/MS application^70,71^. Transitions were summed to obtain isotopologue-level data which was then used to calculate enrichments of each isotopologue on a metabolite level. Enrichments were used for multivariate and rhythmicity analyses. The natural abundance corrected data was also used to calculate phi values (product/precursor relationships) using the Polly Phi LCMS/MS application. The resulting ratios were used for rhythmicity analyses.

The ad libitum vs short-term fasted time course data acquired validated FCF transitions in both ESI negative and positive modes and were used to normalize the data within each mode. After averaging the analytical replicates, a background correction was performed using the Polly Phi LCMS/MS application using the ^12^C injected samples as background samples for the ^13^C injected samples. The background corrected data was then corrected for natural abundance.

### Multivariate and Rhythmicity Analyses

Principal component analyses through SIMCA (Umetrics, version 17) were periodically checked throughout the data processing steps to ensure data quality was being maintained. Enrichments were used for further orthogonal partial least squares discriminant analysis (OPLS-DA) models in SIMCA. OPLS-DA models were generated to test for time dependent within each genotype and/or group by defining each time point as a class with remaining time points as a second class. Genotype/Group level differences were also tested overall as well as for each time point. For each model, the number of components was trimmed to obtain a model with the lowest CV-ANOVA p-value to avoid overfitting. A CV-ANOVA p-value cutoff of 0.1 was used to determine significant models for further analysis. Shared and unique structures (SUS) analyses^72^ were also performed in SIMCA using significant OPLS-DA models. Each metabolite enrichment was treated as an independent variable in multivariate models for computational simplicity.

In addition to multivariate analyses, univariate analyses were also performed. A two-way ANOVA using genotype and time was performed followed by a post-hoc Tukey HSD test for any significant interaction compounds. P-values were adjusted using a Benjamini-Hochberg (BH) correction. The adjusted ANOVA genotype p-values along with calculated fold changes of *fumin* to wild type were used in a volcano plot.

Using the enrichment-level data as 5 biological replicates from ZT 0 to 20, RAIN^73^ and JTK^74,75^ algorithms were used to assess diurnal (24-and 20-28-hr periods) and ultradian (12-hr) rhythmicity through the R Rain and MetaCycle packages, respectively. Results from RAIN using a q-value cutoff of 0.2 were used for further analyses. For the product to precursor relationships (phi values), RAIN p-values less than 0.1 were used to determine significance. For feeding data, rhythms were tested using food consumption combined into 10-minute bins through JTK.

### FCF and Glucose Quantitation

Representative sample of wild type flies were injected with a glucose and blue dye solution, collected, and extracted as described earlier. Uninjected flies were collected and extracted to use as a background matrix for the calibration curve. To ensure a similar background, the upper fractions from uninjected samples were pooled and then aliquoted before drying. Injected samples were resuspended in diluent while uninjected samples were resuspended in appropriate calibration solutions containing varying FCF concentrations in the diluent. Integration was performed in El-Maven as described with data processing in R (version 4.0.3) and Prism (version 9).

### Human Sample Analysis

Plasma samples from 14 patients (8 males and 6 females; 12 Caucasian; average ± standard deviation age 29.14 ± 9.37) collected as described in ^76^ were used. These samples were collected every four hours from an in-patient protocol and the first 24 h of samples analyzed here. The consent of the Institutional Review Board of the University of Pennsylvania and the Clinical and Translational Research Center of the University of Pennsylvania was granted for this study and informed consent obtained from all research subjects. 150 µL of plasma was used as a starting material and extracted using the method described in metabolite extraction and LC-MS measurements section with the following differences: To each sample 900 µL of 2:1 methanol:chloroform was added followed by 300 µL of chloroform and water. Samples were vortexed and centrifuged and 200 µL of the upper fraction was collected for LCMS analyses and dried for 6 hours under vacuum. Paired samples were acquired using an ion-switching method using transitions, solvents, instrumentation, LC and MS parameters described in^38^. For sample resuspension, dried upper fractions were resuspending using 200 µL of 15:85 milliQ water:acetonitrile. Acquired data was integrated using El-MAVEN (version 0.12.1 beta) as described in the data processing and normalization section. Exported ion counts were corrected for instrument drift using a linear correction followed by median fold change normalization. The data was combined for all subjects into a single dataset. To normalize for interindividual differences in concentration, the ratio of each metabolite to the average for that metabolite was taken on an individual level. The resulting ratios were used for multivariate and rhythmicity analyses as described in the multivariate and rhythmicity analysis section.

### Pathway Analysis

Significant pathways as identified by VIP values were used in MetaboAnalyst 5.0 for a pathway analysis^77^. Metabolites were uploaded using HMDB identifiers and processed using a hypergeometric enrichment method, relative-betweenness centrality for topology analysis using the Homo sapiens (KEGG) pathway library. The resulting data was imported into R (Version 4.0.3) and further analyzed. Significant pathways were defined using a FDR less than 0.05. To be included in further analyses, an impact value greater than zero was required for at least one time comparison.

---

**Extended Data Fig. 1:**
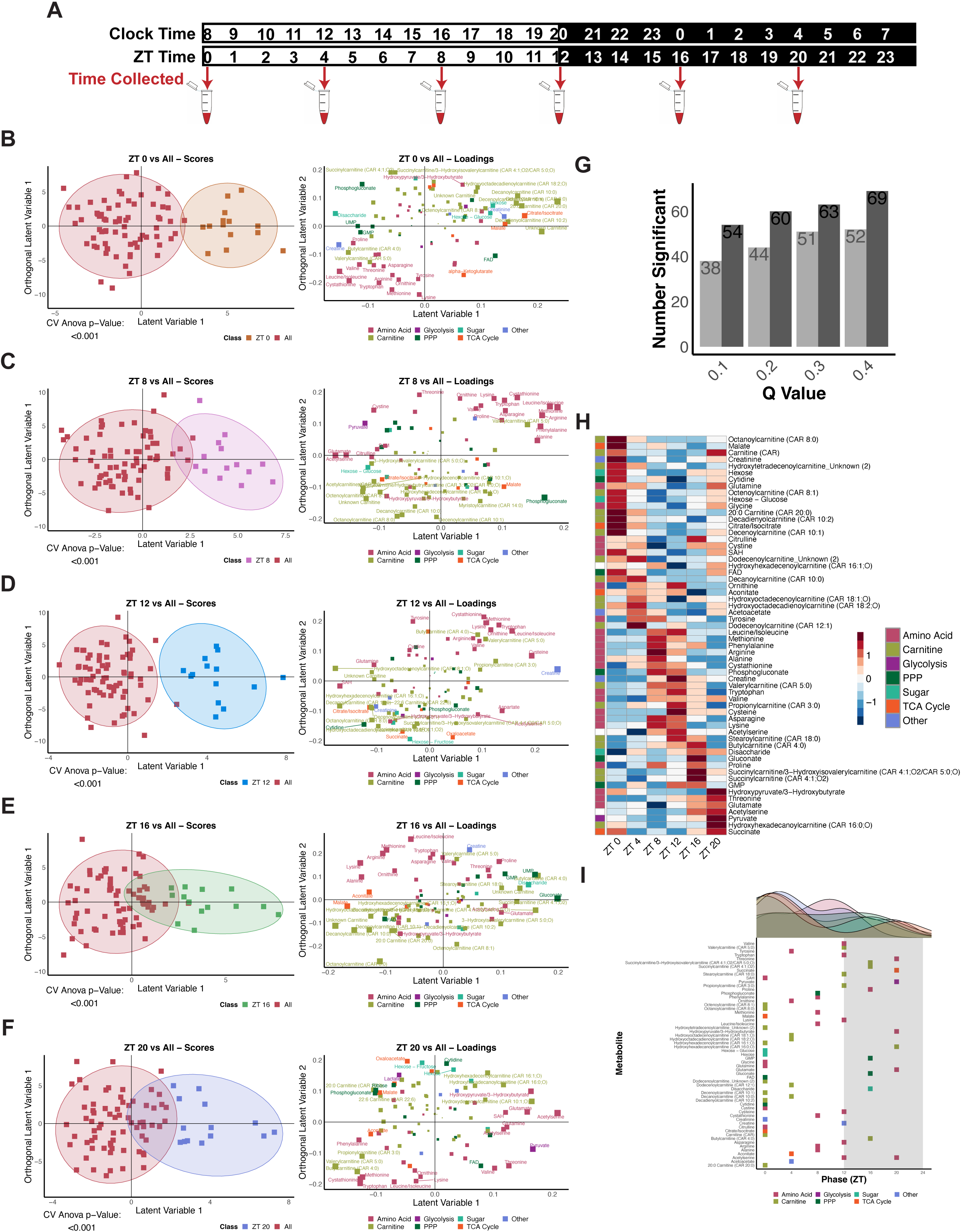
Pairwise OPLS-DA models for each time point and 20–28-hour rhythmicity supports time of day variation in metabolite levels. **a,** Overview of study design showing clock times and corresponding ZT times for blood collections adapted from ^76^. **b-f,** Scores plots (left panel) and corresponding loadings plots (right panel) for significant pairwise OPLS-DA models. Colors in scores plots represent classes and in loadings plot represent classes of metabolites. Size of points in loadings plot represents significance as defined through VIP values. Models are (b) ZT 0 vs All, (c) ZT 8 vs All, (d) ZT 12 vs All, (e) ZT 16 vs All, and (f) ZT 20 vs All. Two classes were defined for each model as the time point being tested and all remaining time points. **g,** Overview of the number of significant 20-28-hr rhythmic metabolites observed at RAIN (dark) or JTK (light) q-value cut-offs of 0.1, 0.2, 0.3, and 0.4. **h,** Phase ordered heatmap of significantly cycling metabolites with 20-28-hr periods as tested by RAIN with a q-value less than 0.2. **i,** Distribution of RAIN phases for significant 20-28-hr metabolites. Colors represent classes of metabolites.

**Extended Data Fig. 2:**
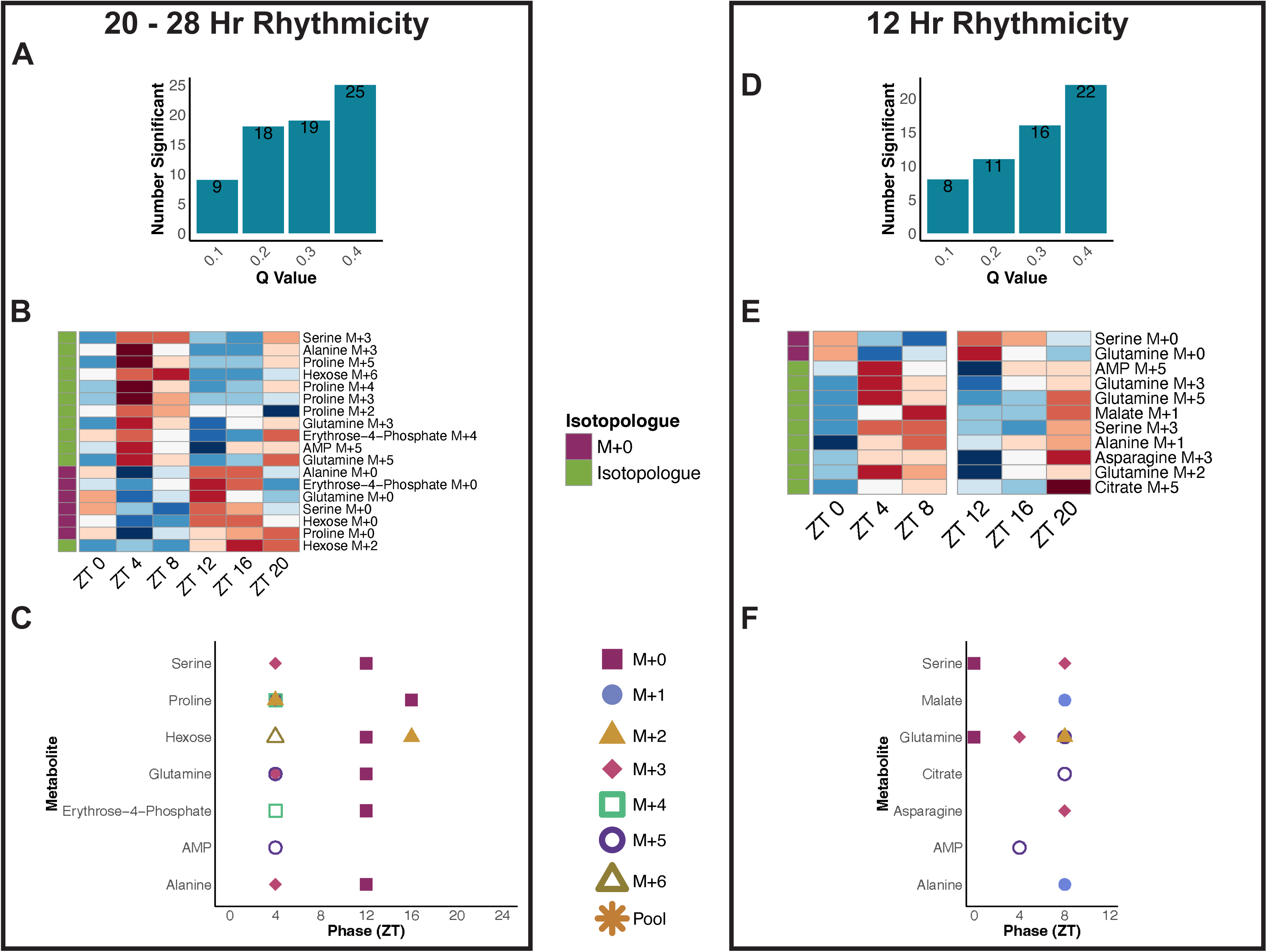
Overview of 20-28-hr and 12-hr rhythmicity in wild type. **a,d,** Overview of the number of significant 12-hr (a) and 20-28-hr (d) rhythmic compounds observed at RAIN q-values of 0.1, 0.2, 0.3, and 0.4. **b,e,** Phase ordered heatmaps of significantly cycling compounds with 12-hr (**b**) and 20-28-hr (**e**) periods in wild type as tested by RAIN with a q-value less than 0.2. **c,f,** Distribution of RAIN phases for significant 12-hr (c) and 20-28-hr (f) compounds grouped by metabolite. Shapes and colors refer to isotopologues or pools.

**Extended Data Fig. 3:**
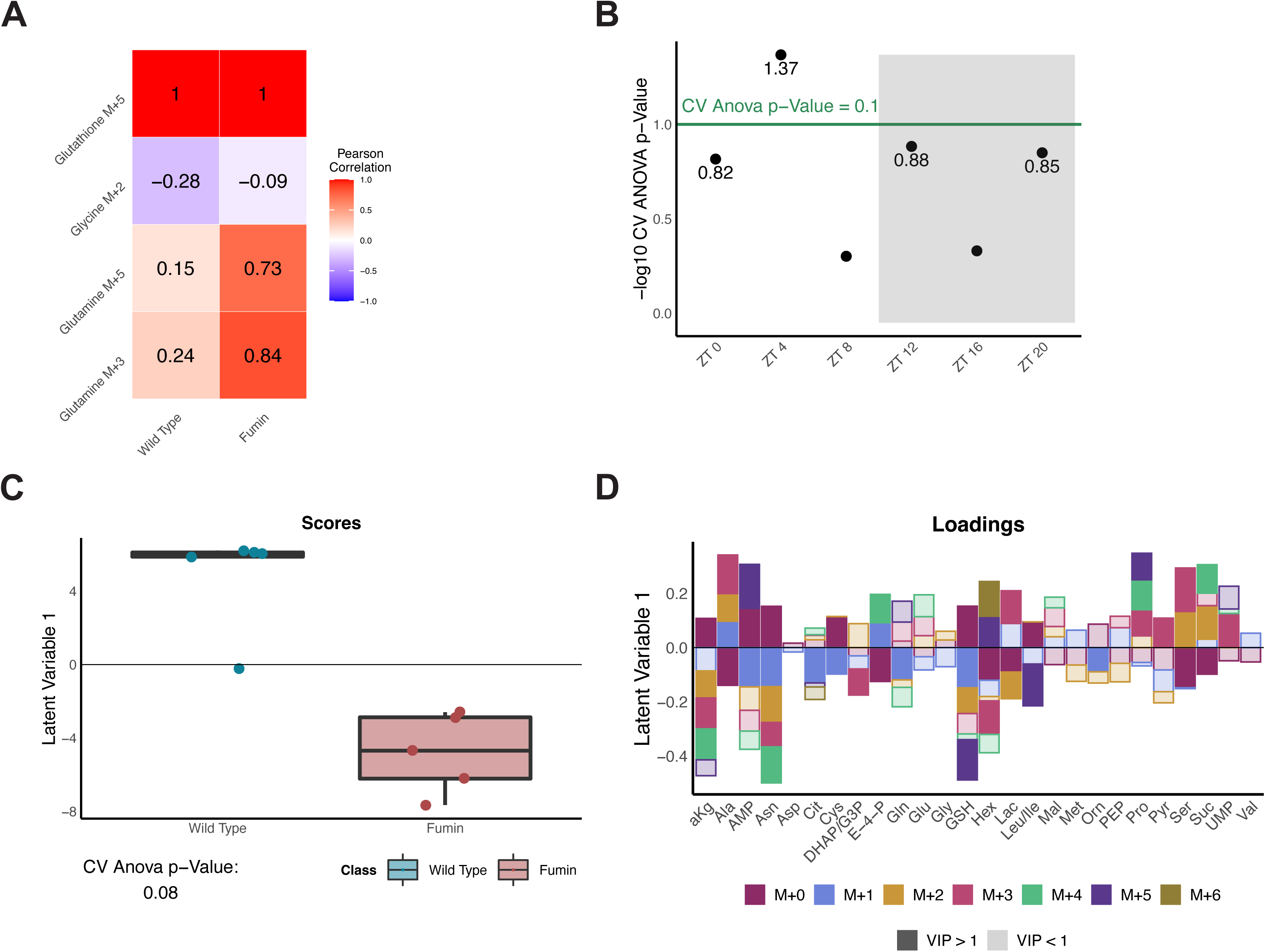
Genotype differences at ZT 4 display common and unique changes in labelled metabolite forms. **a,** Heatmap showing pearson correlation of select isotopologues with glutathione M+5 for both wild type and *fumin*. **b,** Negative log CV-ANOVA p-values for each time point tested in pairwise OPLS-DA models between genotypes. Significance was defined as a CV-ANOVA p-value less than 0.1 (shown as points above the green line). **c,** Scores plot of the significant ZT 4 wild type vs *fumin* OPLS-DA model. Classes were defined as all ZT 4 samples for WT as one class and for *fumin* as a second class. **d,** Loadings plot of the corresponding ZT 4 wild type vs *fumin* OPLS-DA model. Colors represent different isotopologues while shading representing VIP significance (dark signifies a VIP value greater than 1 while lighter shades are used for a VIP less than 1). Abbreviations are defined in Fig. 3g.

**Extended Data Fig. 4:**
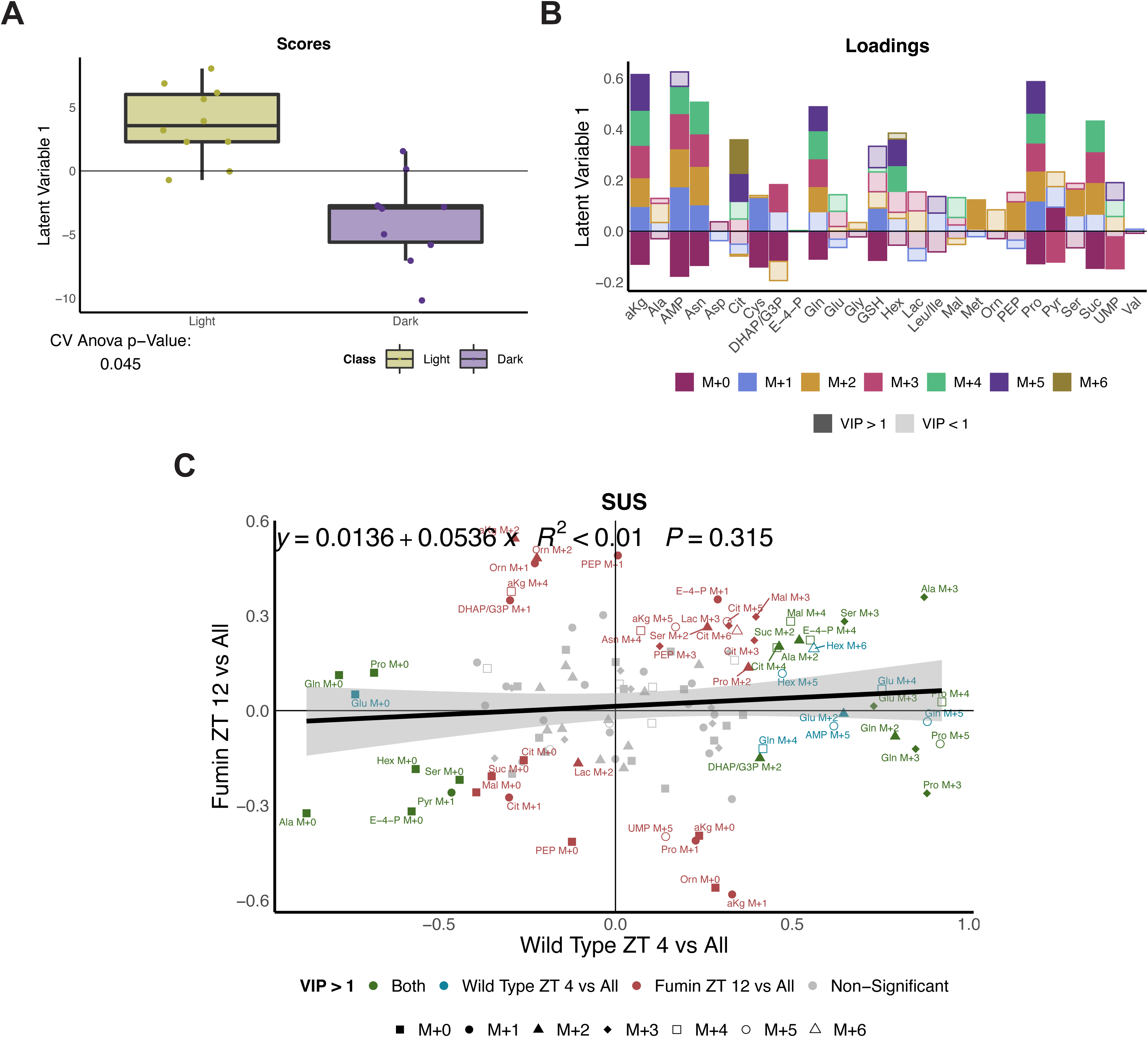
Increased biosynthesis downstream of glucose is observed during the light period in *fumin*. **a,** Scores plot of the significant *fumin* Light (ZT 4 and ZT 8) vs Dark (ZT 16 and ZT 20) OPLS-DA model. **b,** Corresponding loadings plot for the *fumin* Light vs Dark scores (a) plot. Colors represent different isotopologues and shading represents VIP significance. Abbreviations are defined in Fig 3g. **c,** SUS plot comparing the loadings plot from the Wild Type ZT 4 vs All (Fig. 3g) and *fumin* ZT 12 vs All (Fig. 5e) OPLS-DA models with a regression line and equation. Green refers to VIP compounds in both models; blue refers to VIP compounds in the wild type model; red refers to VIP compounds in the *fumin* model. Shapes refer to isotopologues. Abbreviations are defined in Fig. 3g.

**Extended Data Fig. 5:**
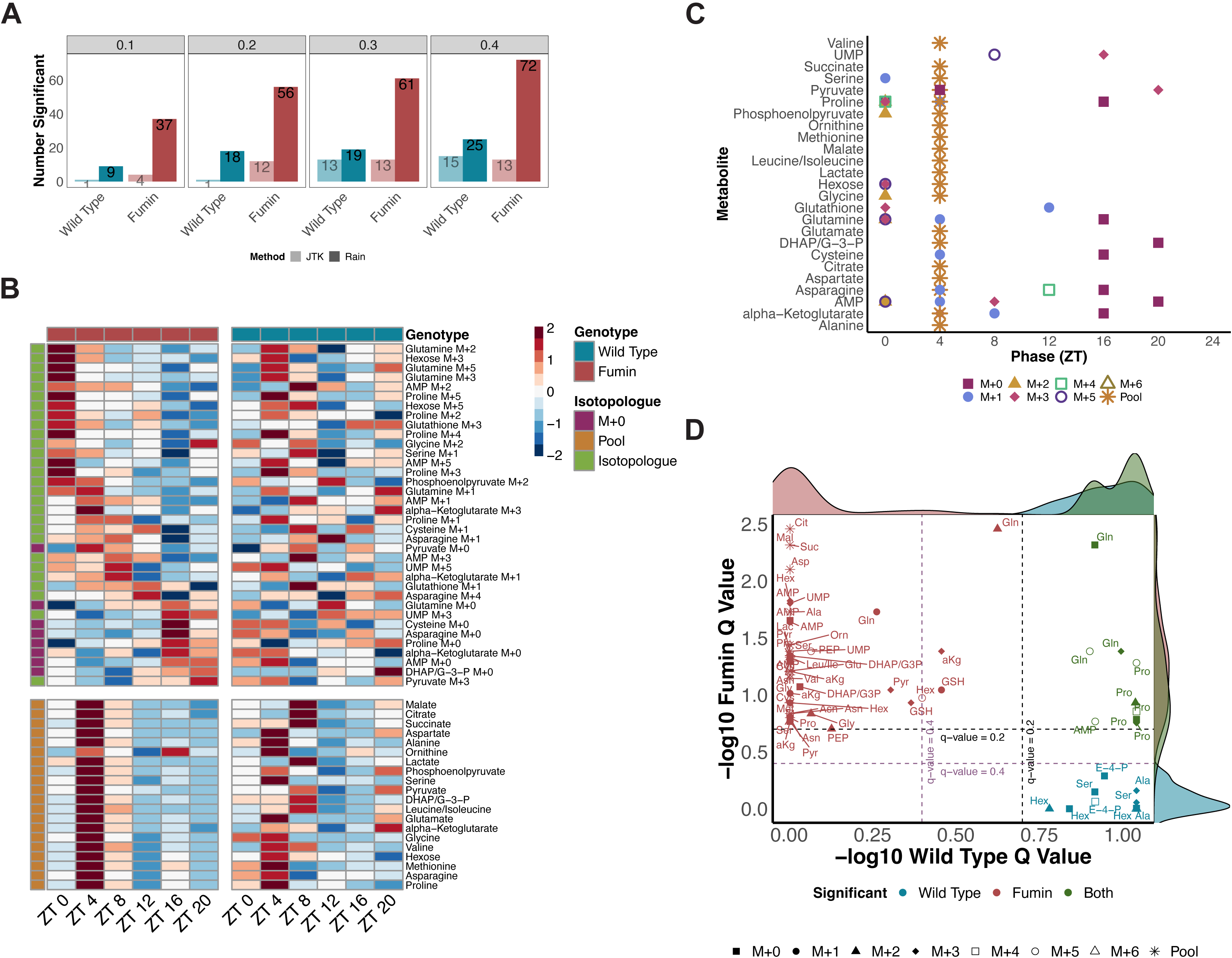
20-28-hr rhythmicity in *fumin* complements/supports 24-hr rhythmicity. **a,** Overview of the number of significant 20-28-hr rhythmic compounds observed at RAIN (dark) or JTK (light) q-value cut-offs of 0.1, 0.2, 0.3, and 0.4. **b,** Phase ordered heatmaps of significantly cycling compounds with periods of 20-28-hr in *fumin* at a RAIN q-value cutoff of 0.2. Enrichments for wild type are shown for comparison. **c,** Distribution of RAIN phases for significant 20-28-hr compounds in *fumin* grouped by metabolite. Shapes and colors refer to isotopologues or pools. **d,** Comparison of 20-28-hr significant cycling compounds in wild type and *fumin* as defined by a RAIN q-value less than 0.2. q-value cutoffs of 0.2 (black dotted line) and 0.4 (purple dotted line) are shown. Color refers to significant rhythmicity in one or both genotypes while shapes refer to isotopologues. Abbreviations are defined in Fig. 3g.

**Extended Data Fig. 6:**
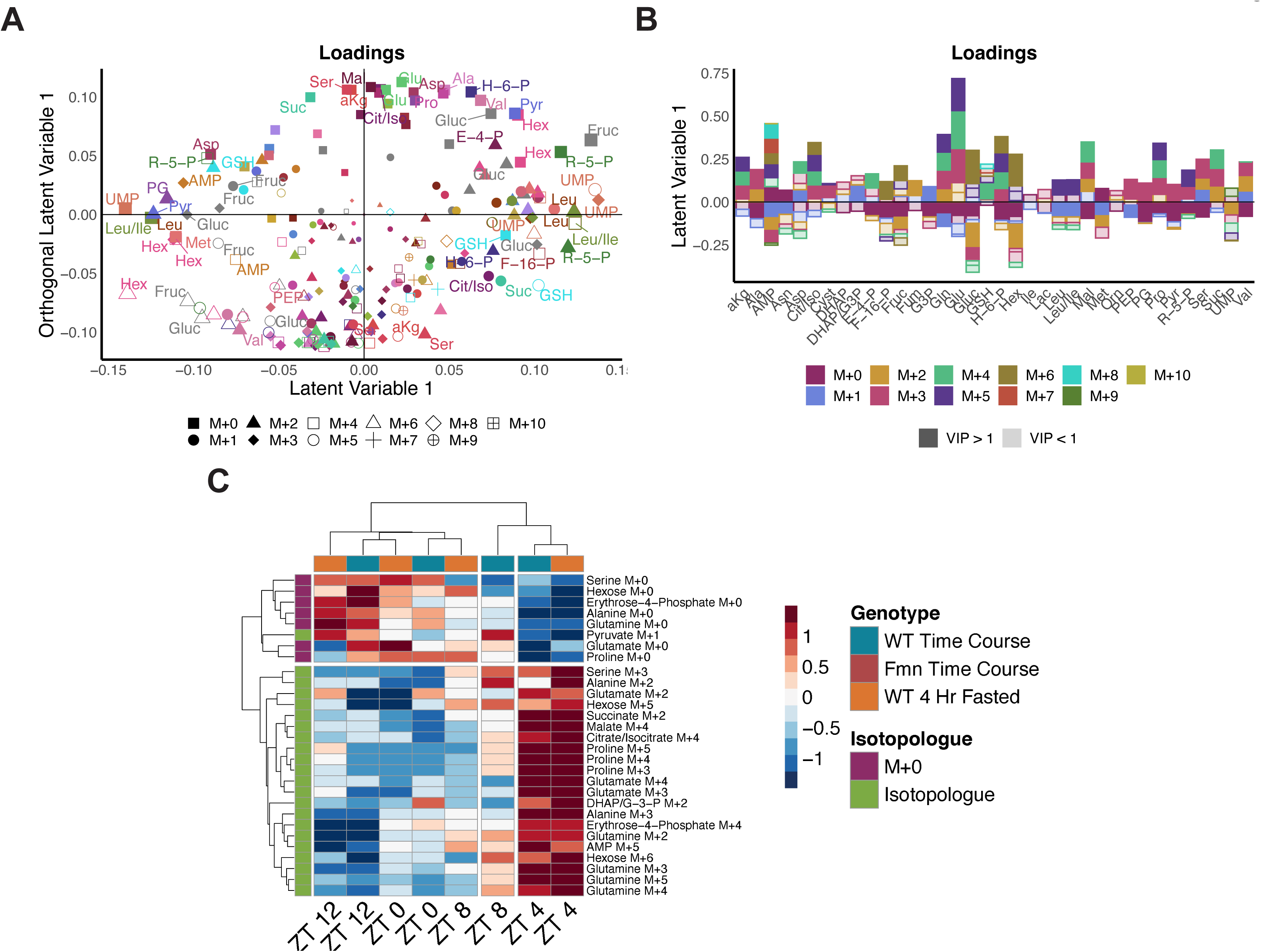
Changes in isotopologue enrichments in response to a 4-hr short term fast. **a,** Corresponding loadings plot for the ad libitum vs short-term fasted condition. Isotopologues are represented by shapes, metabolites by color and size of the shapes represents significance as defined through VIP values. Abbreviations as defined in Fig. 3g; Isocitrate (Iso); Fructose-1,6-Phosphate (F-1,6-P); Fructose (Fruc); Fumarate (Fum); Glucose (Gluc); Hexose-6-Phosphate (H-6-P); Phosphoglycerates (PG); Ribulose-5-Phosphate (R-5-P). **b,** Corresponding loadings plot for the short-term fasted ZT 4 vs All OPLS-DA model. Colors represent isotopologues with shading representing VIP significance (dark signifies a VIP value greater than 1 while lighter shades are used for a VIP less than 1). Abbreviations as defined in Fig. 3g and a. **c,** Clustered heatmap on both the x and y-axes shown using ZT 0 to 12 time points from the earlier Wild Type samples along with the short-term fasted samples for VIP compounds determined from the Wild Type ZT 4 vs All model (Fig. 3f and g).

## Supplemental Tables

Extended Data Table 1: Transitions for wild type and *fumin* time courses

Extended Data Table 2: Transitions for wild type *ad libitum* and 4-hour fasted time courses

Extended Data Table 3: Positive mode raw data for wild type and *fumin* time courses

Extended Data Table 4: Negative mode raw data for wild type and *fumin* time courses

Extended Data Table 5: Enrichment and pool values for wild type and *fumin* time courses

Extended Data Table 6: Positive mode raw data for wild type *ad libitum* and 4 hour fasted time courses

Extended Data Table 7: Negative mode raw data for wild type *ad libitum* and 4 hour fasted time courses

Extended Data Table 8: Enrichment and pool values for wild type *ad libitum* and 4-hour fasted time courses

## Highlights

Fly micro capillary injections allow for high time resolution metabolic tracing Biosynthesis peaks in the early light phase from a labelled glucose challenge Biosynthesis rhythms are enhanced in a hyperactive dopamine transport mutant Rhythms in biosynthesis are robust to four-hour food restriction or activity

